# Targeted LNPs deliver mRNA encoding IL-15 superagonists to balance efficacy and toxicity in cancer therapy

**DOI:** 10.1101/2024.01.11.575299

**Authors:** Juntao Yu, Qian Li, Shenggen Luo, Xiaona Wang, Qiang Cheng, Rongkuan Hu

## Abstract

Interleukin-15 (IL-15) emerges as a promising immunotherapeutic candidate in oncology because of its pivotal role in modulating both innate and adaptive immunity. However, the therapeutic utility remains concern due to the unexpected toxicity. We propose here that the mRNA lipid nanoparticle (mRNA-LNP) system can balance the issue through targeted delivery to increase IL-15 concentration in the tumor area and reduce leakage into the circulation. Utilizing the Structure-driven TARgeting (STAR) platform, we acquired intellectual property LNP vectors for effective and selective mRNA delivery to local (LNP^Local^) and to pulmonary (LNP^Lung^). Then the promising IL-15 superagonists mRNAs were obtained through structural optimization and sequence screening, showing better activity compared with benchmarker N-803. Subsequently, the anti-tumor efficacy of IL-15 superagonists mRNAs were evaluated by intratumoural (i.t.) injection and intravenous (i.v.) injection via LNP^Local^ and LNP^Lung^, respectively. As a result, such superagonists exhibited better anti-tumor activity, less systematic exposure, and less cytokine related risks than N-803. We finally verified the selective delivery and well tolerability of LNP^Lung^ in non-human primates (NHPs), confirming the potential for clinical application. This finding may open up new possibilities for the treatment of lung cancers and lung metastasis cancers.

## Introduction

The critical role that Interleukin-15 (IL-15) plays in innate and adaptive immunity has had profound implications in the field of immunology (*1, 2*). The biological effects of IL-15 are mediated through interaction with a receptor complex comprising three distinct chains: CD122 (IL-2/IL-15Rβ), CD132 (IL-2/IL-15Rγc), and the high-affinity-specific chain IL-15Rα (*3*). IL-15 currently has garnered considerable attention as a promising immunotherapeutic candidate for cancer treatment (*4, 5*), exemplified by the advancement of the IL-15 analogue and agonist complex agent nogapendekin alfa (N-803) (*6–9*), as well as the ongoing clinical trials of similar agents (*10–12*). However, potential toxicity remains due to the mechanism by which IL-15 superagonists activate T/NK cells (*13–15*). Such potential toxicity was believed from the shared activities of IL-15 and IL-2, such as the increase in IL-15 observed in autoimmune disorders (*16*), and detection of high level of IL-15/IL-15Rα positively associated with leukemia (*17–20*). During treatment, therapeutic enhancement of CD8 T cells, NK cells, and in some cases CD4 T cells was observed at all doses and routes of administration of IL-15 (*21, 22*). Of note, higher doses and more frequent administrations have resulted in more pronounced responses but also cause clinical toxicities, such as anorexia, diarrhea, weight loss, and transient grade 3-4 neutropenia (*23–26*). Therefore, the initial indication for the N-803 New Drug Application (NDA) submitted for the treatment of bladder cancer was based on intravesical administration (*27*), which can relatively balance therapeutic efficacy and potential toxicity compared with systemic injection. Likewise, we believe that this balance could also be achieved through targeted delivery to achieve maximum IL-15 concentration in the target area but reduce leakage into the circulation.

The mRNA technology may become one of the options for above goal via targeted delivery and expression. The success of mRNA technology in COVID-19 vaccines has propelled it into a global research focus, marking a rapid advancement (*28*). Through sequence design, mRNA can encode a wide range of proteins, offering extensive application prospects (*29–32*). Delivered through lipid nanoparticles (LNP), mRNA technology has been successfully applied to the defense and treatment of multiple diseases, making this delivery platform one of the most important factors in clinical translation (*33–38*). However, targeted delivery of mRNA-LNP to specific organs and cells remains challenging, which could be partially satisfied by high-throughput screening, component optimization, and antibody modification (*34, 39–41*). Through rational design, we have synthesized hundreds of ionizable cationic lipids and established a Structure-driven TARgeting (STAR) platform, in which tissue-selective mRNA-LNP was fully validated (PCT/CN2023/137998). Notably, LNP^Local^ and LNP^Lung^ from STAR LNPs library could specifically deliver mRNA into muscle and lung, respectively. Therefore, these two LNPs may meet the needs of tumor therapy by delivering mRNA encoding IL-15 superagonists, especially in designed tissues.

Herein, we firstly evaluated the structural stability and activity of several IL-15 superagonists in the form of mRNA. Among them, the STR-4 mRNA, with a structure of immunoglobulin G Fc fragment (Fc) – IL-15Rα sushi domain – IL-15 design, showed the best efficacy that was even better with established benchmark N-803. Followed, the sequence-optimized mRNAs (STR-4-D43 and STR-4-D61, both derived from STR-4 sequence by introducing aspartic acid mutation in specific sites) were performed for anti-tumor experiments in subcutaneous tumor model and lung metastasis tumor models. Delivered by LNP^Local^ via i.t. injection and LNP^Lung^ via i.v. injection, both mRNAs showed significant tumor inhibition. Moreover, the mRNA-LNP modality showed well-balanced efficacy and safety via minimizing IL-15 leakage to major organs, reducing unexpected toxicity. In order to test the potential of clinical translation, we finally verified the robust delivery efficiency and well-safety of mRNA-LNP^Lung^ on non-human primate (NHP). These findings suggest that targeting mRNA-LNPs may provide another promising option for the IL-15 based cancer treatment, especially in lung cancers or lung metastatic tumors.

## Results

### Validation of targeted mRNA-LNPs

To improve the therapeutic window of IL-15, we propose that mRNA-LNP system may satisfy the goal via targeted delivery. On the one hand, the structure and sequence of IL-15 superagonists can be refined and optimized in the form of mRNA to enhance their anti-tumor activity. Targeted LNPs, on the other hand, could satisfy limited higher concentrations of IL-15 around the tumor area but minimize leakage into the circulation to reduce unintended side effects (**Fig. 1A**). We previously have prepared a library of ionizable cationic lipids through rational design and patented the LNP platform, named Structure-driven TARgeting (STAR). Notably, two distinct LNPs from STAR LNP library demonstrated the high efficacy and specificity for muscle (LNP^Local^) and lung (LNP^Lung^) mRNA delivery (**Fig. 1B**). The particle size of LNP^Local^ was 84.93 ± 2.72 nm, and that of LNP^Lung^ was 53.23 ± 1.36 nm, both with uniform polydispersity index (PDI) (**Fig. 1C**). The Cryo-TEM images revealed a highly consistent shape for both LNP^Local^ and LNP^Lung^ samples (**Fig. 1, D and E**). The encapsulation efficiency (EE) for LNP^Local^ and LNP^Lung^ was determined to be 94.72 ± 3.58 and 99.77 ± 0.05, respectively. Furthermore, stability testing indicated negligible changes in particle size and PDI for both LNPs after 28 days of storage at 4°C (**Fig. 1, F and G**), affirming their substantial stability and suitability for *in vivo* investigations. Through intramuscular (i.m.) injection, mRNA-LNP^Local^ exhibited enhanced efficiency and reduced hepatic leakage compared with positive SM102 LNP the widely-validated mRNA vector (**Fig. 1H**). The quantification analysis revealed that the percentage of muscle expression was 97.61% for LNP^Local^ and 88.69% for the SM102 LNP (**Fig. 1I**). These results convincingly demonstrated that LNP^Local^ enhanced local targeting and reduced hepatic extravasation after i.m. injection (**Fig. 1, I and J**). Similarly, through intravenous (i.v.) delivery, the expression of firefly luciferase (Fluc) was identified in the lungs by mRNA-LNP^Lung^ (**Fig. 1, K and L**). Bioluminescence in the lungs accounted for 98.18% of overall bioluminescence, with negligible leakage (0.71%) in the liver (**Fig. 1M**). Such result underscores the remarkable ability of LNP^Lung^ to specifically target the lungs. Overall, the results indicated that both LNP^Local^ and LNP^Lung^ had the potential to specifically deliver IL-15 superagonist mRNA and broaden the therapeutic window.

**Fig. 1.**
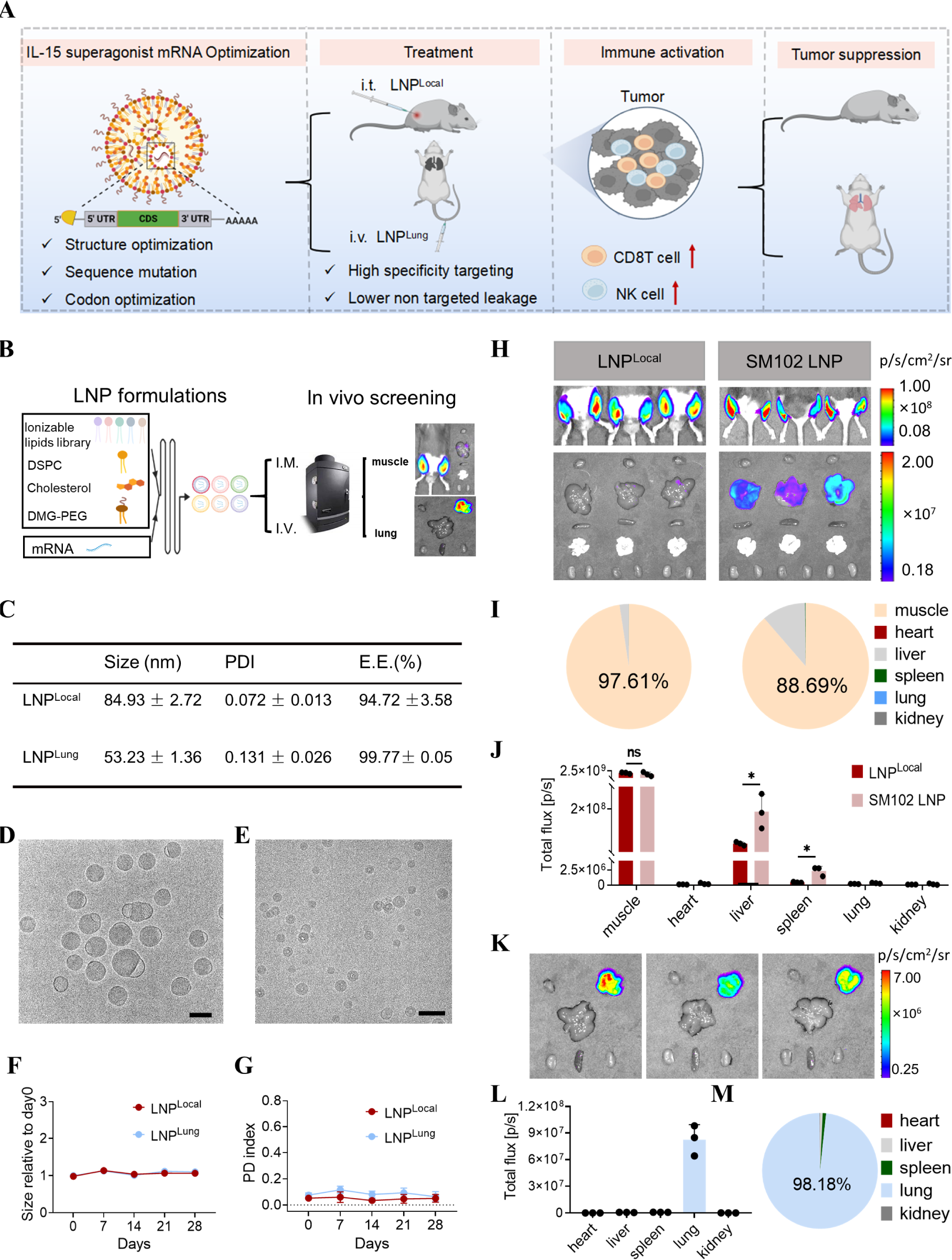
Evaluation of targeted LNP formulations. (**A**) Schematic illustration of IL15 mRNA-LNP balancing safety and efficacy in cancer therapy. (**B**) Schematic illustration of established Structure-driven TARgeting (STAR) platform, in which mRNA-LNP could reach muscles and lung specificity. (**C**) Characterization of LNP^Local^ and LNP^Lung^ encapsulating firefly luciferase (Fluc) mRNA. (**D, E**) Images of Cryo-electron microscopy of LNP^Local^ and LNP^Lung^ (Scale bar = 100 nm). (**F, G**) Size and PDI change over 28 days at 4℃. (**H-M**) Validation of previously designed LNPs through the Fluc expression *in vivo*. (**H**) IVIS images of LNP^Local^ and SM102 LNP for delivering Fluc mRNA via i.m. injection (0.05 mg/kg/leg). (**I**) Percentage of luciferase expression in muscles and various organs of LNP^Local^ and SM102 LNP. (**J**) Total flux (ROI) of luciferase signals measured in muscles and other organs treated by various mRNA-LNPs. (**K**) IVIS images of LNP^Lung^ for delivering Fluc mRNA by i.v. injection (0.25 mg/kg). (**L**) Total flux (ROI) of luciferase signals measured in different organs after injection of LNP^Lung^. (**M**) Percentage of luciferase expression in various organs treated by mRNA-LNP^Lung^. Data is shown as mean ± s.e.m. (n = 3 biologically independent samples or animals).

### The structural optimization of mRNA encoding IL-15 superagonist

The sushi domain of IL-15Rα has been recognized for stabilizing IL-15(*42*), by which the N-803 was progressed to NDA submission in the Food and Drug Administration (FDA) in 2022. Despite this achievement, the structural organization of the IL-15-IL-15Rα complex remains unexplored. To address this, we established a classification system comprising four distinct categories for IL-15-IL-15Rα (sushi) complexes (**Fig. 2A**). Among them, STR-1 and STR-2 represent direct fusion of IL-15 to IL-15Rα (sushi) on N-terminus and C-terminus, respectively. STR-3 and STR-4 involve the addition of the immunoglobulin G (IgG) Fc fragment to STR-1 and STR-2, potentially improving pharmacokinetic parameters. *In vitro* analyses of protein-level assay revealed that STR-2 and STR-4 (unbound C-terminal IL-15) exhibited reduced EC_50_ values than N-803 in IL-2Rβ-STAT5 signaling system (**Fig. 2B**). Considering IL-2Rβ-STAT5 signaling is the key signaling pathway of IL-15 downstream activation in its target cell – lymphocytes (*43*), this result indicated that STR-2 and STR-4 might have better potency than N-803. This claim was supported by subsequent data derived from peripheral blood mononuclear cells (PBMCs) of lung cancer patients. After co-cultured with mRNA expressed STR-2 and STR-4 supernatant, a significantly elevated level of pSTAT5 and interferon gamma (IFNγ) was observed in lung cancer patients’ PBMCs co-culture system, compared to samples co-cultured with N-803 (**Fig. 2, C and D**). In addition, in a co-culture system with PBMCs from healthy donors, all mRNA direct-expressed supernatant group presented a significant elevated proliferation rate than N-803 group (**Fig. 2E**), indicating an excellent activity enhancement of mRNA modality. Moreover, the proliferation of target cells (CD8+ T and CD56+ NK cells) was observed in all treated groups, with STR-2 and STR-4 appearing to be better (**Fig. 2F**). These findings suggested that structural optimization of IL-15 superagonist was critical, and unbound C-terminal IL-15 design was more attractive.

**Fig. 2.**
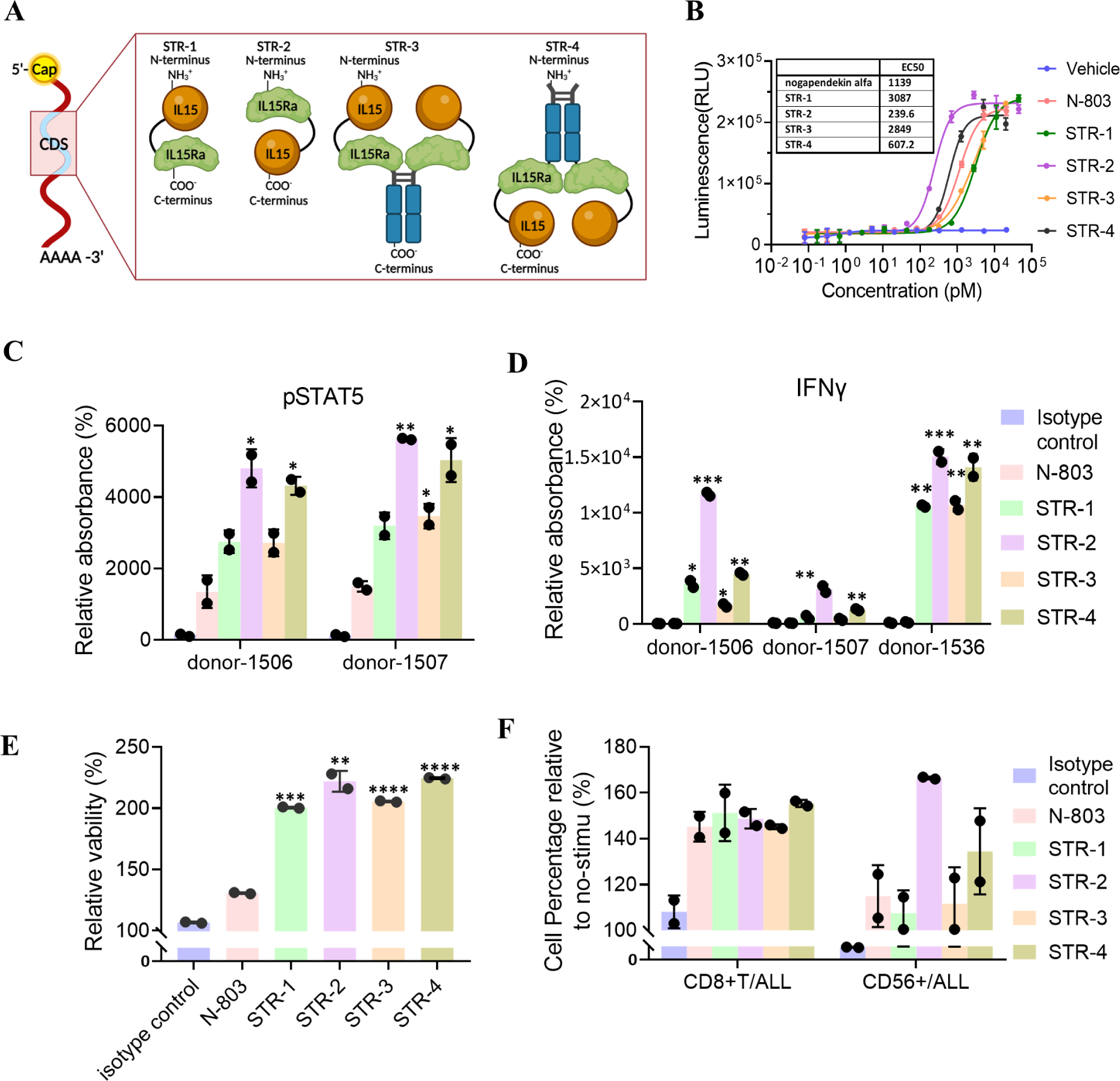
Improvement of *in vitro* activity of mRNA-encoded IL-15 superagonists. **(A)** The schematic illustration of structural optimization of the IL-15 superagonist via mRNA format. **(B)** Detection of activating signal of IL2Rβ-STAT5 in TF-1 IL2Rβ-STAT5 reporter cells. Reporter cells were co-cultured with cell medium containing either N-803 or IL-15 superagonist protein for 24 h before detection. **(C-F)** Assessment of the activation of peripheral blood mononuclear cells (PBMC) in a co-culture system with N-803 or mRNA-encoded IL-15 superagonists supernatant. PBMC were co-cultured with N-803 (4.5 nM) or mRNA-encoded IL-15 superagonists supernatant (0.1 nM), as well as 10 ug/ml CD3Ab (OKT3) for 3 days, then several parameters were detected, including (**C**) pSTAT5 levels in the cell lysate, (**D**) IFNγ levels in the supernatant, (**E**) cell proliferation rate, and (**F**) changes in CD45+CD3+CD8+ and CD45+CD3+CD56+ cell subtype proportion. The PBMC samples were collected from lung cancer patients (**C, D**) and healthy individuals (**E, F**). Data is shown as mean ± SEM and asterisks indicate significant differences between the N-803 and mRNA-encoded IL-15 superagonist groups (*, P < 0.05; **, P < 0.01; ***, P < 0.001; ****, P < 0.0001).

### The anti-tumor efficacy of STR-2 and STR-4 in subcutaneous tumor models

To further test the anti-tumor activity between STR-2 and STR-4, we applied mRNA-LNP^Local^ delivery system in subcutaneous heterotopic tumor models (**Fig. 3A**). Through intratumoral (i.t.) injection, STR-2 and STR-4 exhibited similar efficacy in inhibiting tumor growth compared to the group treated with N-803 in the CT26 tumor model, and the anti-tumor effect was dose-dependent (**Fig. 3B, D and table S1**). Importantly, the N-803 group displayed significant reduction in body weight, indicating systemic distress and significant sickness, whereas the groups treated with mRNA-encoded IL-15 superagonists (STR-2 and STR-4) maintained consistent body weight throughout the study (**Fig. 3C**). It is worth mentioning that the dosage of mRNA-LNP (2 mg/kg) was twice that of N-803 (1 mg/kg), indicating that the mRNA-LNP^Local^ encoded IL-15 superagonists could well-balance the efficacy and *in vivo* toxicity. Pharmacokinetics studies demonstrated that N-803 group (0.2 mg/kg) had higher drug concentrations in the plasma and larger area under the curve (AUC) compared to the 10 times higher dose mRNA-LNP groups (2 mg/kg) (**Fig. 3E**), which was believed to resulted in toxicity associated with systemic exposure to IL-15 superagonists. Then the superior anti-tumor efficacy of STR-2 and STR-4 were further supported in the B16F10 tumor model (**Fig. 3, F and G**). Considering the toxicity of N-803, a dose of 0.2 mg/kg is used here. As expected, no significant decrease in body weight was observed in the 10-fold higher dose (2 mg/kg) of the mRNA groups (**Fig. 3H**). Additionally, the STR-4 showed more pronounced efficacy in inhibiting tumor growth compared to STR-2 and N-803 (**Fig. 3, G and I, table S2**). Therefore, we selected STR-4 in the followed processes.

**Fig. 3.**
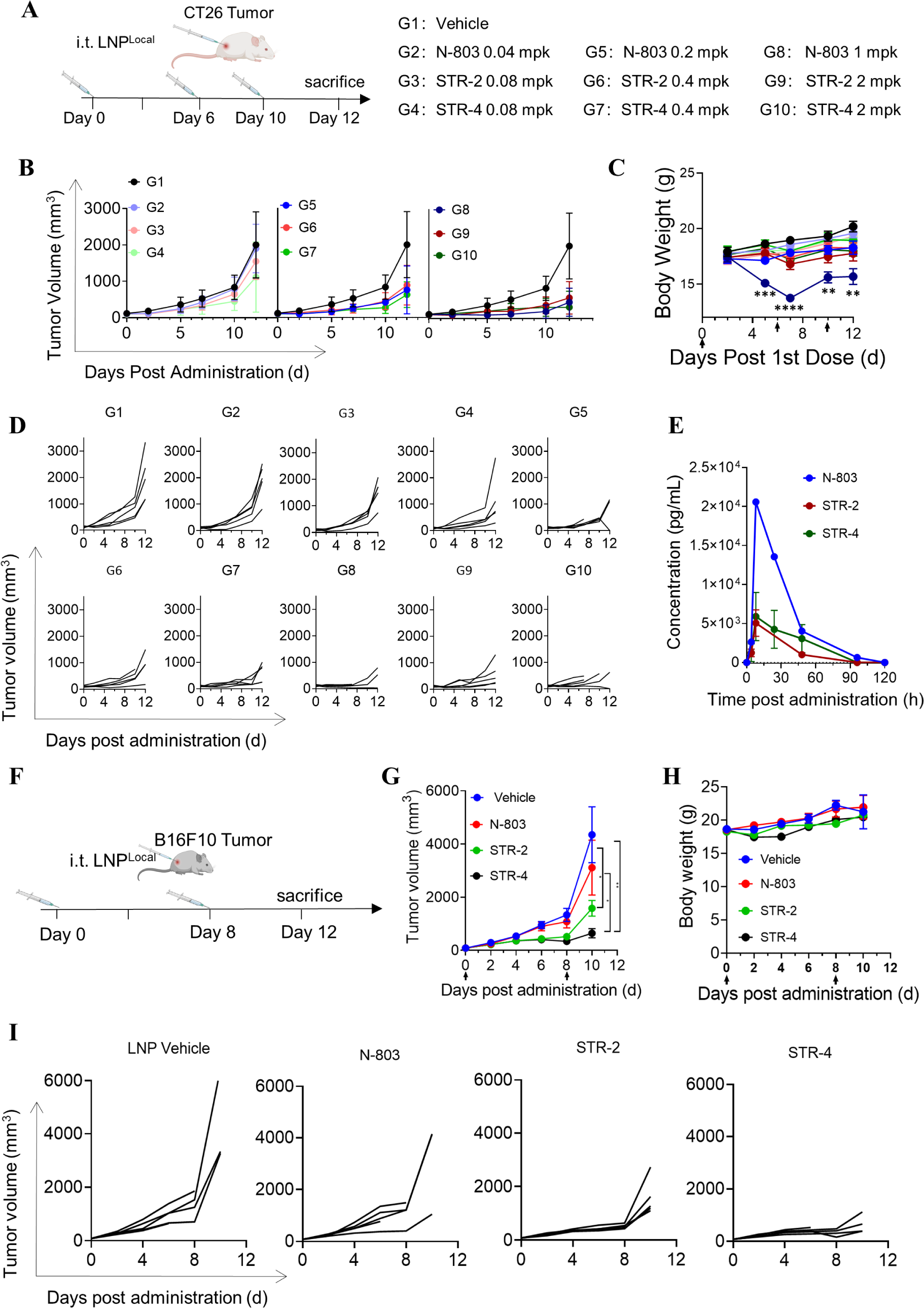
The mRNA-LNPs encoding STR-2 and STR-4 well-balanced efficacy and toxicity in subcutaneously tumor model. **(A)** The schematic diagram of optimized STR-2 and STR-4 mRNA for anti-tumor assay delivered using LNP^Local^. Mice were inoculated subcutaneously with 1E6 CT26 tumor cells and randomized into groups when tumors reached 80-120mm^3^, then mice were dosed three times and marked day 0, day 6 and day 10. Each group, N-803 (i.v.), SRT-2 (i.t.) and SRT-4 (i.t.) included low, medium and high doses. **(B, D)** Tumor growth in each group was monitored cross day 0 to day 12. **(C)** The change of body weight was recorded from the day post 1^st^ dose. **(E)** The pharmacokinetics of the agents in mice, with plasma concentrations measured before dosing and at 4h, 8h, 24h, 48h, 72h, 96h, and 120h after a single dosing. **(F)** The schematic diagram of optimized STR-2 and STR-4 mRNA for anti-tumor assay delivered using LNP^Local^ in B16F10 model. Mice were inoculated subcutaneously with 1E6 B16F10 cells and randomized into groups when tumors reached 80-120mm^3^. Then the mice were treated by various groups, vehicle (empty LNP, i.t.), N-803 (0.2 mpk, i.v.), SRT-2 (2 mpk, i.t.) and SRT-4 (2 mpk, i.t.), for twice (marked day 0 and day 8). The tumor growth (**G, I**) and body weight changes (**H**) were recorded since the day post dosing to day 12. Data is shown as mean ± s.e.m. (n = 5 biologically independent animals). Asterisks indicate significant differences (*, P < 0.05; **, P < 0.01; ***, P < 0.001; ****, P < 0.0001).

### Sequence optimization of mRNA encoding STR-4

Previously study reported that mutating amino acid (AA) into aspartic acid (Asp, D) in IL-15 agents could potentially improve its receptor binding affinity and downstream signaling activity (*44*), which inspired us to further optimize the sequence of STR-4 for seeking better efficacy. We first performed codon optimization on STR-4 and obtained STR-4-D0 (without D mutation). A series of mutated STR-4 sequences were then generated by one-by-one “to D” AA mutation from the N-terminus to the C-terminus in the IL-15 portion, resulting in 108 different mutants **(Fig. 4A)**. For example, STR-4-D61 indicates that the 61^st^ not-D-AA of the IL-15 portion is mutated to D. Then the 108 mutants were evaluated to initiate IL-2Rβ-STAT5 signaling in reporter cells (**Fig. 4B**). Five of them (STR-4-D43, STR-4-D56, STR-4-D61, STR-4-D65 and STR-4-D106) showed strong ability (**Fig. 4C, fig. S1, and table S3**). Furthermore, all the five variants showed similar or better binding affinity to IL-2Rβ/γ complex compared to STR-4-D0 (**fig. S2**), indicating a potential ‘enrichment to target’ capability. All them demonstrated better activity than N-803 in inducing proliferation of CD8+ T cells and NK cells subclasses in PBMCs co-culture system, with the STR-4-D61 showed the best (**Fig. 4, D and E**). We also noticed that STR-4-D43 showed only moderate activity in reporter cells and PBMC *in vitro* activation, but the highest expression level in the HEK293T (**fig. S3**), suggesting that it may be able to satisfy the anti-tumor activity at lower dosage of mRNA for potentially reducing a formulation related toxicity.

**Fig. 4.**
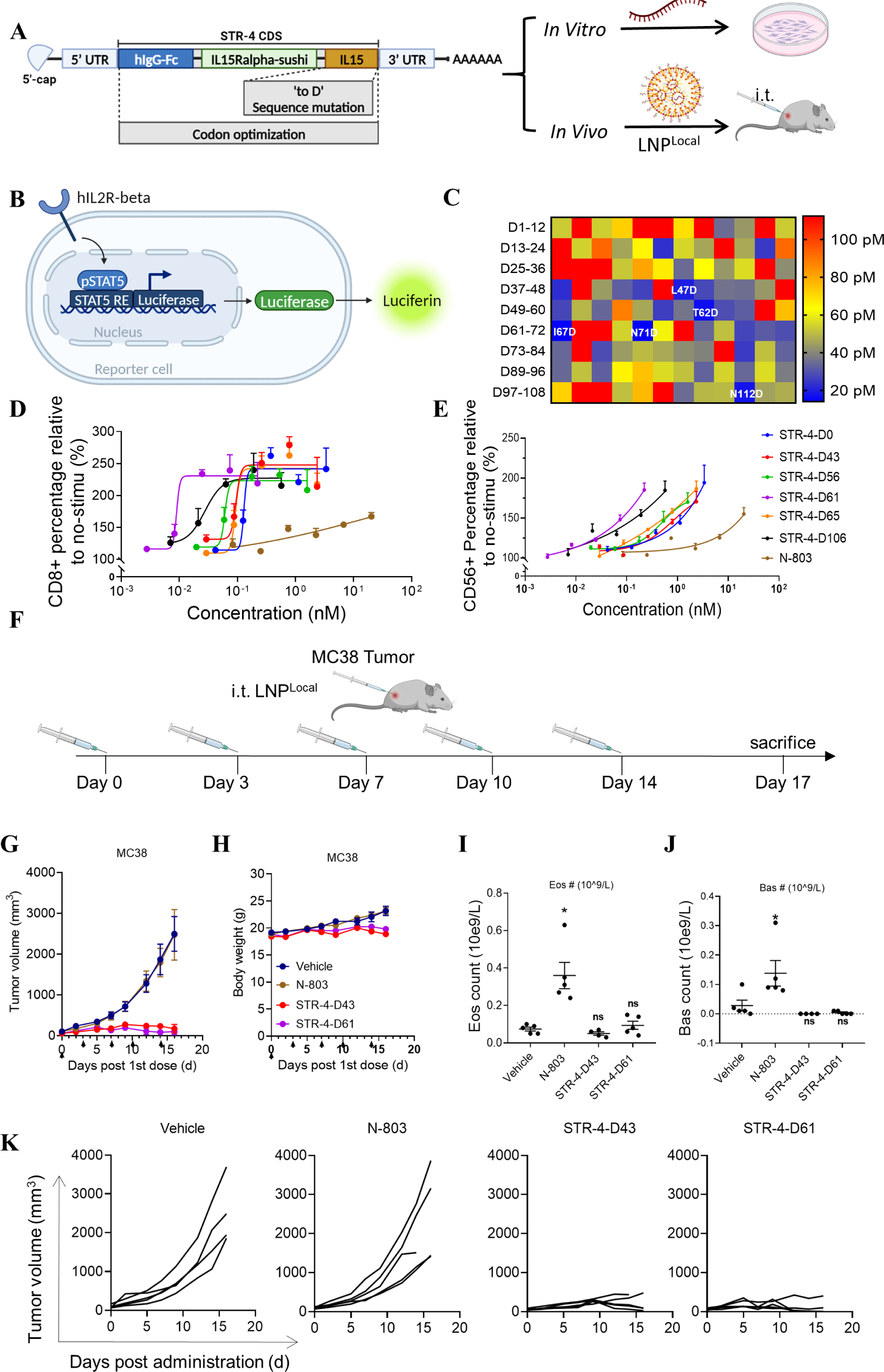
Sequence optimization of STR-4. **(A)** This schematic illustrates the design and process of STR-4 mRNA sequence mutation, codon optimization, and *in vitro* and *in vivo* screening. **(B)** The schematic diagram of assessing the ability to activate IL2Rβ-STAT5 signaling in TF-1 reporter cells. **(C)** The EC_50_ heat map of 108 different STR-4 mutants, which was generated by one-by-one “to D” AA mutation from the N-terminus to the C-terminus in the IL-15 portion. **(D**, **E)** Assessment of the activation of peripheral blood mononuclear cells (PBMC) in a co-culture system with top STR-4 mutants. The changes in CD45+CD3+CD8+ **(D)** and CD45+CD3+CD56+ **(E)** cell subtypes were analyzed, respectively. **(F)** The schematic diagram of STR-4-D43 and STR-4-D61 mRNAs for anti-tumor assay delivered by LNP^Local^ in MC38 model. Mice were inoculated subcutaneously with 5E5 MC38 cells and randomized into groups when tumors reached 80-120mm^3^. Then mice were treated by various groups, vehicle (empty LNP, i.t.), N-803 (0.2 mpk, i.v.), SRT-4-D43 (2 mpk, i.t.) and SRT-4-D61 (2 mpk, i.t.), at day 0, day 3, day 7, day 10, and day14. Tumor growth **(G**, **K)** and body weight changes (**H**) were monitored during whole experiment, and **(I**, **J)** blood samples were collected on day 17 for Complete Blood Count. Data is shown as mean ± s.e.m. (n = 3-5 biologically independent samples or animals). Asterisks indicate significant differences (*, P < 0.05; **, P < 0.01; ***, P < 0.001; ****, P < 0.0001).

We thus tested the *in vivo* anti-tumor activity of STR-4-D61 and STR-4-D43 delivered by LNP^Local^. The experiments were performed using subcutaneously transplanted MC38 tumor model in C57BL/6 mice. When tumors reached a volume range of 80-120 mm^3^, the mice were randomly grouped and i.t. administered on specific days (day 0, day 3, day 7, day 10, and day 14) (**Fig. 4F**). Across groups, N-803 showed limited effectiveness in inhibiting tumor growth, whereas the mRNA-encoding groups showed significant antitumor activity (**Fig. 4, G and K**). Importantly, no statistically reduction of body weight was observed, indicating the well-tolerated property (**Fig. 4H**). To illustrate the superiority of mRNA-LNP modality more clearly, we analyzed several blood indicators in tumor-bearing mice. Although N-803 treatment presented confined anti-tumor activity, the increasing level of the total number of white blood cells, IL-6 levels, eosinophils, and basophil subpopulations in the N-803 group were significantly observed (**Fig. 4I-J, fig. S4, and fig. S5**). Considering lymphocytes contribute to most of the IL-15 related anti-tumor activity (*45*) and IL-6 was one of the key factor to cytokine related toxicity (*46, 47*), these phenomena may explain why mRNA-LNP showed lower toxicity *in vivo*. Furthermore, the levels of red blood cells, hemoglobin, and platelets in the mRNA-LNP groups were all detected within the normal range (**fig. S5**), verifying the hematological safety of this strategy.

### Anti-tumor efficacy of STR-4-D61 and STR-4-D43 in the lung metastatic model

To further expand the antitumor potential of mRNA-LNP approach, we intend to test the possibility of systemic delivery, which is very challenging due to uncontrollable toxicity of IL-15 related agents mentioned above. From the STAR LNP library, we used LNP^Lung^ and LNP^Liver^ to deliver STR-4-D61 mRNA and detected the target protein concentrations after i.v. injection. Six hours post injection, a very high concentration (∼ 2×10^5^ pg/mL) of related protein was detected in the plasma from STR-4-D61 LNP^Liver^ group, and then the concentration was maintained over 100 h. As contrast, the protein concentration was below the limit of detection (50 pg/mL) 24h after injection in the plasma of STR-D61 LNP^Lung^ group, and the peak concentration was very low (∼2×10^3^ pg/mL) compared to LNP^Liver^ group (**Fig. 5A**). This result demonstrated that STR-4-D61 LNP^Lung^ may be able to balance the safety and efficacy in treatment of lung cancer or lung metastasis by reducing IL-15 leakage. To verify this, anti-tumor activity of STR-4-D61 LNP^Lung^ was employed in B16F10 lung metastatic tumor models. The C57BL/6 mice were firstly i.v. injected with luciferase-expressed B16F10 cells, then were treated by STR-4-D61 LNP^Lung^ at indicated days (**Fig. 5B**). At given time-points, mice were imaged by *in vivo* imaging system (IVIS). Compared with control group, STR-4-D61 LNP^Lung^ showed notable reduction or complete elimination of luciferase expression (**Fig. 5, C and D**). This was also confirmed by isolated lung tissue images, in which lots of tumor lesions were observed in the vehicle group (**Fig. 5E**). Moreover, similar experiment was performed with STR-4-D43 LNP^Lung^ as well (**Fig. 5F**). STR-4-D43 LNP^Lung^ treated mice also exhibited remarkable tumor inhibition detected by IVIS (**Fig. 5, G-I**). The lung sections of hematoxylin and eosin (H&E) staining confirmed the anti-tumor efficacy that STR-4-D43 LNP^Lung^ group showed a close-to-healthy lung tissue (**Fig. 5K**). We further analyzed the numbers of cytotoxic T cells and natural killer (NK) cells in blood. The proportion of CD161+ (mouse NK cell marker) cells were observed obviously in STR-4-D43 LNP^Lung^ group compared to vehicle group (**Fig. 5J**). Considering NK cell is one of the key target cells of IL-15 superagonist and play key roles in tumor killing, this result indicated that the mRNA-LNP^lung^ system can successfully activate its target. We also noticed the obvious lung infiltration of CD3+T cell and CD8+ T cell in STR-4-D43 group (**Fig. 5L**), which was a remarkable signal of anti-tumor immunity activation. All of these results illustrated that STR-4 variants LNP^lung^ achieved outstanding anti-tumor efficacy in lung metastasis tumor model by mobilization of local CD8+T cells and NK cells.

**Fig. 5.**
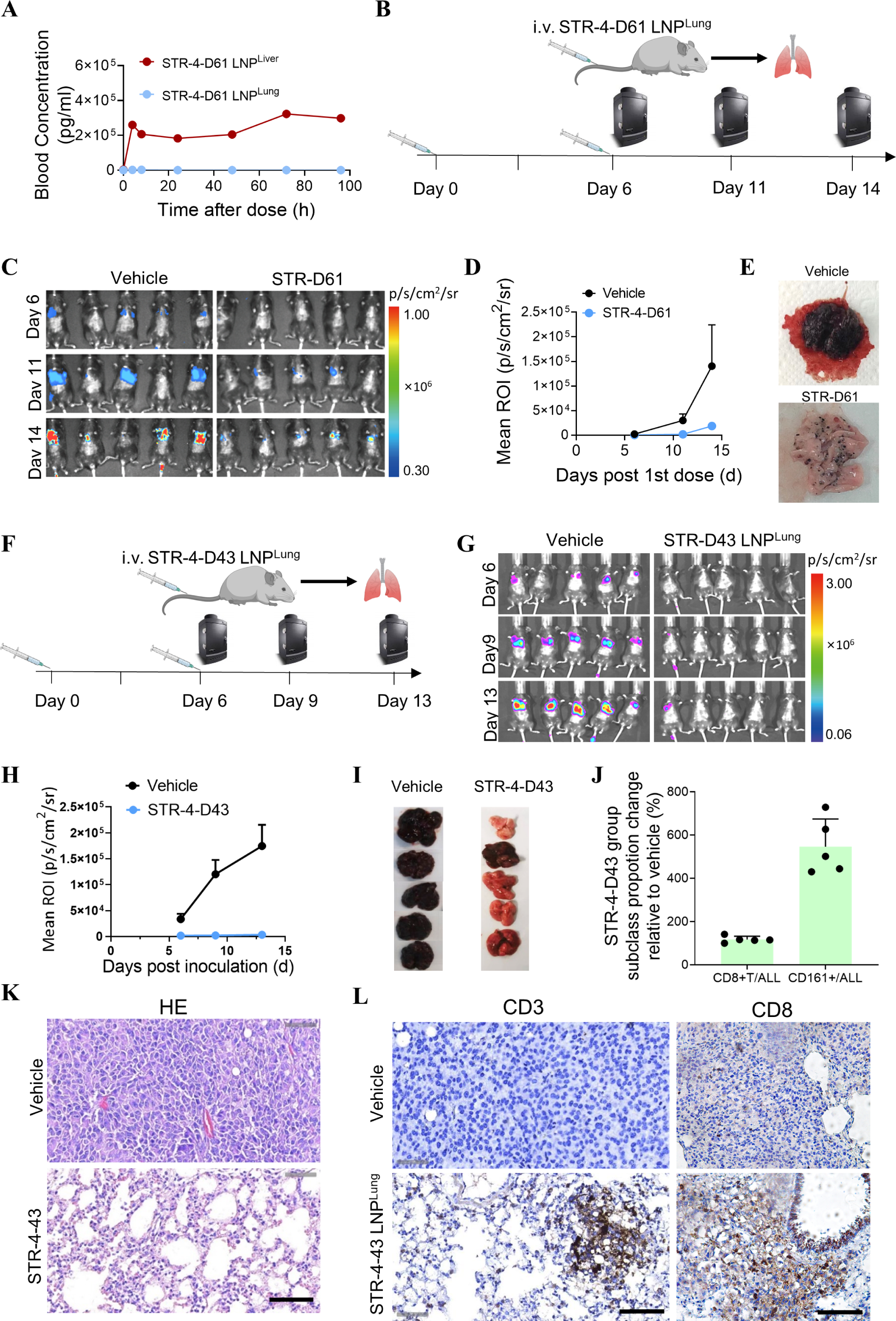
The remarkable anti-tumor activity of top STR-4 variants in lung metastatic tumor. **(A)** The comparison of STR-4-D61 protein in blood after the mRNA was delivered with LNP^liver^ and LNP^lung^ by i.v. injection. **(B)** The schematic diagram of STR-4-D61 for lung metastatic tumor therapy. Mice were inoculated intravenously with luciferase-expressing B16F10 and randomized into groups. After 6h, mice were i.v. treated by STR-4-D61 LNP^Lung^ for twice with mRNA dose of 1 mg/kg. **(C, D)** Tumor growth of each group was monitored by IVIS system and quantification data was analyzed. **(E)** At day 14, mice were sacrificed and lung images were recorded. **(F)** The antitumor effect of STR-4-D43 mRNA on lung metastatic tumors was also evaluated through experiments similar to those described above. **(G-I)** Obvious tumor inhibition of STR-4-D43 group was observed by IVIS imaging, luciferase quantification and lung tissue photographs. **(J, L)** Treated by STR-4-D43 LNP, the proportion of CD161+ NK cells in blood were significantly increased, and the increase infiltration of CD3+T cells and CD8+ T cells in the lung was observed. (**K**) H&E staining confirmed the anti-tumor efficacy that STR-4-D43 group showed a close-to-healthy lung tissue. Scale bar = 100 μm. Data is shown as mean ± s.e.m. (n = 5 biologically independent animals).

### Targeted delivery and safety evaluation of LNP^Lung^ in NHP

To further evaluate the potential of clinical application using mRNA STAR LNP modality, a few experiments were conducted in non-human primate (NHP). We selected mRNA-LNP^Lung^ systems with i.v. administration into Macaca fascicularis, in which one female was treated by 1xPBS, another female and a male were treated by Fluc mRNA-LNP^Lung^ (**Fig. 6A**). Two doses of injections were completed at 0 h (0.25 mg/kg empty LNP) and 24 h (0.125 mg/kg mRNA-LNP), respectively. After another 6 h, NHPs were sacrificed and major tissues were imaged by IVIS. Compared with the control group, very strong luciferase expression in lungs were detected in both NHPs with almost undetectable expression in the liver and spleen. The expression percent of the lungs reached 94.46% and 86.90%, respectively (**Fig. 6B-D**). Immunohistochemistry (IHC) analysis also confirmed that Fluc was expressed in the lungs, with significantly higher compared to the control group (**Fig. 6E**). These results fully demonstrated the efficiency and specificity of mRNA-LNP^lung^ in NHPs. At the same time, we assessed the safety by analyzing several blood parameters, including albumin (ALB), blood urea nitrogen (BUN), urea (UA), and lymphocytes (LYM). All indicators are within normal range, no obvious difference compared with control group was observed (**Fig. 6F-I**), indicating excellent safety feathers of mRNA-LNP^lung^ in liver, kidney and immune system. Overall, these results demonstrated that LNP^Lung^ exhibited a promising lung targeting efficacy and safety in NHPs, resulting great potentials for clinical translation.

**Fig. 6.**
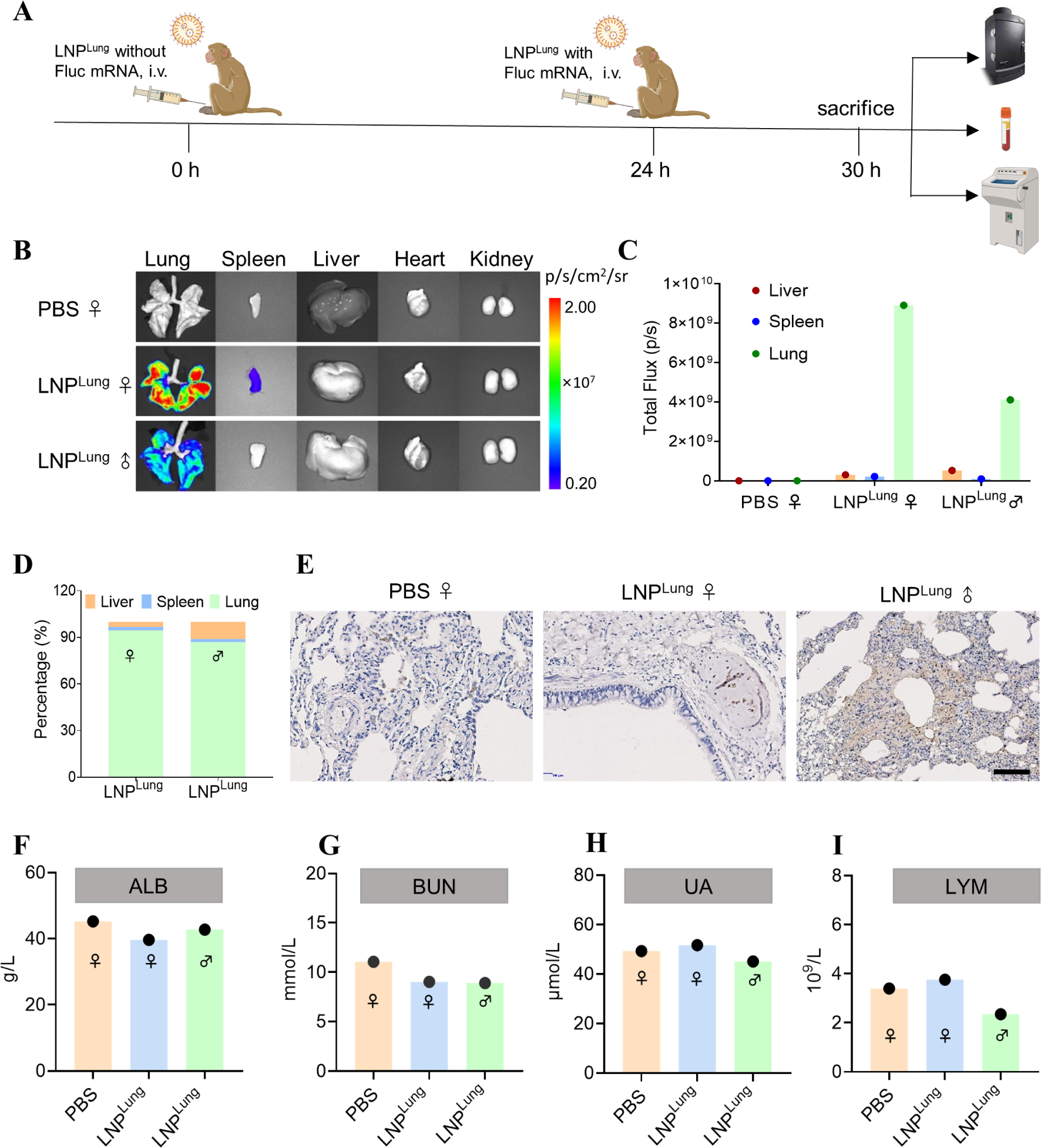
The validation of efficacy and safety of mRNA-LNP^Lung^ in non-human primates (NHPs). (**A**) Schematic illustration of design of experiment in Macaca fasciculiris. One female was treated by 1xPBS, another female and a male were treated by Fluc mRNA-LNP^Lung^. Two doses of injections were completed at 0 h (0.25 mg/kg empty LNP) and 24 h (0.125 mg/kg mRNA-LNP), respectively. After another 6h, tissues and blood were collected for further analysis. (**B-D**) Luciferase expression of major tissues was imaged, quantified and analyzed. (**E**) Immunohistochemistry (IHC) images were performed to confirm the Luc expression in the lungs (Scale bar, 100 μm). (**F-I**) The blood parameters were assessed to verify the safety of mRNA-LNP^Lung^. ALB: albumin, BUN: blood urea nitrogen, UA: urea, LYM: lymphocyte.

## Discussion

As the National Cancer Institute’s immunotherapy workshop in 2007 recognized the potential to revolutionize cancer treatment, the IL-15 has been widely acknowledged. Multiple clinical studies have been exploring the feasibility of IL-15 related agents as a potential anti-tumor therapy (*48, 49*). However, its dual effects from T/NK cell interaction raise concerns about on-target toxicity, leading to inadvertent infiltration and attacks on non-tumor organs (*50, 51*). This issue arises from the activation of effector T cells and natural killer (NK) cells, whose cytokine production can exacerbate these effects (*51*). Therefore, IL-15-based systemic therapies face challenges for dosage limitation during treatment. A well-known IL-15 superagonist, N-803, thereby selected local perfusion when submitted the initial indication for NDA for bladder cancer treatment (*27*).

Considering these evidences, it is reasonable that maximizing IL-15 concentration in the tumor region and minimizing leakage into circulation may provide an attractive strategy for cancer treatment. In this work, the proof-of-concept (POC) experiments were performed based on our established mRNA-LNP platform (named STAR LNP) via targeted delivery of IL-15 superagonist mRNA for cancer therapy.

We firstly optimized the structures of IL-15 superagonists and demonstrated that C-terminal IL-15 structure with prolonged pharmacokinetics feature, known as STR-4, exhibited the best efficacy. Then we evaluated the STR-4 variants by introducing D mutation in the domain of IL-15, and screened out STR-4-D61 and STR-4-D43 mRNAs which showed even better activity. Regarding the targeted delivery, both LNP^Local^ and LNP^Lung^ could effectively and specifically deliver mRNA into targeted area and significantly decreased the leakage into circulation, which may benefit the balance of efficacy and safety for IL-15 based cancer therapy. The followed anti-tumor experiments showed that mRNA-LNP modality indeed presented better therapeutic effects but negligible toxicity compared with positive N-803, even at 2-10 fold higher doses. More importantly, we verified the delivery efficacy, specificity, and safety of mRNA-LNP^Lung^ in NHP level, which may accelerate the clinical translation on lung cancer or lung metastasis cancer therapy.

Overall, this work provides an attractive strategy for IL-15-based cancer therapy by targeted mRNA-LNP platform. This technology can balance efficacy and safety and is expected to expand the therapeutic window of IL-15. It is worth mentioning that the STAR LNP platform also includes those LNPs that can achieve mRNA delivery specificity to the liver, brain, etc., so we have reason to believe that the reported method is also able to treat the cancers there. Still, more experiments are needed for further improvements. For example, this work only analyzed luciferase expression and preliminary toxicity in NHP levels, which may not be sufficient to give very solid conclusions. Additional tests including multiple mRNA expression, time-dependent expression, acute and long-term toxicity assessment are needed and will be discussed in the future.

## Materials and Methods

### Materials

For LNP preparation. SM102 、 DSPC 、 Cholesterol and DMG-PEG were purchased from SINOPEG (China). The ionizable cationic lipids used in LNP^Lung^ and LNP^Liver^ were from Starna Therapeutics Co., Ltd.. Pur-A-Lyzer Midi Dialysis Kits (WMCO, 3.5 kDa) were purchased from Sigma. The Quant-iT^TM^ RiboGreen^TM^ RNA Assay Kit was obtained from Invitrogen (USA). For mRNA preparation. The T7 High Yield Transcription Kit was purchased from ThermoFisher (MA, USA). The Quant-iT^TM^ RiboGreen^TM^ RNA Assay Kit was obtained from Invitrogen (USA). The mRNA was purified using the Monarch RNA Cleanup Kit (NEB, MA, UK). Followed reagents were used for IL-15 superagonist characterization. Human IL-15 Quantikine ELISA Kit (R&D Systems, MN, USA), 96-well ELISA plates (Corning, NY, USA), recombinant human IL-15Rbeta/gamma Heterodimer Protein (Sino Biological, Beijing, China), N-803 (Sino Biological, Beijing, China), Anti-Human IgG-Fc Secondary Antibody (Sino Biological, Beijing, China), ELISA basic kit (Multi Science, Hangzhou, China), One-Lite Luciferase Assay System (Vazyme, Nanjing, China), ImmunoCult™-XF T Cell Expansion Medium (StemCell Technologies, Vancouver, Canada), CD3Ab OKT3 (Sino Biological, Beijing, China), Human IFN-gamma Quantikine ELISA Kit (R&D systems, MN, USA), Human IL-6 Quantikine ELISA Kit (R&D systems, MN, USA), Mouse PBMC Isolation Kit (Solarbio, Beijing, China), CellTiter-Glo® Luminescent Cell Viability Assay (Promega, WI, USA), STAT5 alpha/beta (Phospho) [pY694/pY699] Human InstantOne™ ELISA Kit (ThermoFisher, MA, USA), anti-mouse CD3 antibody (Proteintech, Rosemont, USA), anti-mouse CD8 antibody (Abclona, Woburn, USA) and anti-mouse CD161 antibody (Abcam, Cambridge, UK). Antibodies listed below. FACS related antibodies (anti-human CD45-APC-Cy7 antibody, anti-human CD3-FITC antibody, anti-human CD8-BV510 antibody, and anti-human CD56-PE-Cy7 antibody, anti-mouse CD45-APC-Cy7 antibody, anti-mouse CD3-FITC antibody, anti-mouse CD8-PE-Cy7 antibody, and anti-mouse CD161-APC antibody, and cell fix/perm solutions) were all purchased from BD Pharmingen (NJ, USA). Immunohistochemistry related antibodies include CD3, CD8 and CD56. The CD3 antibody was purchased from Protentech, CD8 antibody was gained from Abclona, and the CD56 antibody was obtained from Abcam. The secondary antibody used for immunohistochemistry was purchased from BOSTER. For cell lines. HEK293T cells (Chinese Academy of Sciences Cell Bank, Beijing, China), B16F10 cells (Cobioer Biosciences, Nanjing, China), MC38 cells (Cobioer Biosciences, Nanjing, China), and CT26 cells (Cobioer Biosciences, Nanjing, China) were cultured in Dulbecco’s Modified Eagle Medium (manufactured by HyClone, UT, USA), supplemented with 10% FBS (Gibco, MA, USA), and 1% Penicillin-Streptomycin (Gibco, USA). IL-15Rbeta-STAT5 reporter TF-1 cells (Genomeditech, Shanghai, China) were cultured in Roswell Park Memorial Institute (RPMI) 1640 medium supplemented with 10% FBS, 1% Penicillin-Streptomycin, and 2 ng/ml GM-CSF (Sino Biological, Beijing, China). Human peripheral blood mononuclear cells (PBMCs) were sourced from OriBiotech (Shanghai, China) and cultured in ImmunoCult™-XF T Cell Expansion Medium (StemCell Technologies, Vancouver, Canada). Mouse PBMCs were purified from mouse whole blood by Mouse PBMC Isolation Kit (Solarbio, Beijing, China). All the mentioned cell lines were maintained under sterile conditions and cultured at 37°C in a 5% CO2 environment.

### LNP formulation

The construction of mRNA-loaded lipid nanoparticle (LNP) formulations was accomplished by microfluidic mixing using a previously described method (*39, 41*). Briefly, The SM102 LNP formulation was prepared with a molar ratio of SM102:DSPC:Chol:DMG-PEG = 50/10/38.5/1.5. The formulations for LNP^Local^ and LNP^Lung^ were developed based on our patented methodology. Specifically, LNP^Local^ formulation was prepared with a molar ratio of ionizable lipid:DSPC:Chol:DMG-PEG = 46/10/42.4/1.6, LNP^Liver^ formulation was same as the LNP^Local^, LNP^Lung^ formulation was prepared with a molar ratio of ionizable lipid:DSPC:Chol:DMG-PEG = 70/4/23.9/2.1. In this process, all lipid components with specific molar ratios were dissolved in ethanol, while the mRNA was dissolved in a 10 mM citrate buffer at pH 4.0. The synthesis procedure employed microfluidics, utilizing an ethanol to water phase volume ratio of 1/3 and a flow rate of 1/3. Subsequently, a dialysis step was conducted in 1×PBS for a duration of 2 hours.

### mRNA design and synthesis

The expressing validation mRNA luciferase vector was constructed by luciferase natural coding sequence (CDS). The design of IL-15 superagonists adhered to the specifications outlined in Figure 2A. The construction of related vectors was conducted by Genescript (Nanjing, China). The specific mRNA sequence was generated through *in vitro* transcription, employing the T7 High Yield Transcription Kit. Following transcription, the mRNA was purified using the Monarch RNA Cleanup Kit. This process ensured the production of purified mRNA for subsequent experiments.

### Characterization of LNP

The size and polydispersity index of LNPs were assessed using dynamic light scattering (BECKMAN COULTER DelsaMax PRO, or Malvern Panalytical Zetasizer Pro). The mRNA encapsulation efficiency was calculated according to the instruction of the Quant-iT™ RiboGreen™ RNA Assay Kit. The Cryo-electron microscopy was taken by the Electron Microscope Platform of Shanghai Jiao Tong University, Zhang Jiang Institute for Advanced Study. To study LNP stability, the size and polydispersity index were monitored for 4 weeks during storage in PBS at 4℃.

### Animal welfare

All mice used in this study were housed and handled in accordance with the Animal Welfare Act and the Regulations Relating to the Use of Animals in Research. All mice related studies followed the principles of the ARRIVE guidelines to ensure the quality and transparency of animal experiments. The number of animals used in this study was minimized by statistical methods, and all possible measures were taken to reduce the pain and distress of the animals during the experiment. The mice study presented in Figure 1 was approved by the Institutional Animal Care and Use Committee (IACUC) of the Peking University Animal Center Ethics Committee. Other mice studies were approved by the IACUC of Kaiji Pharmaceutical Technology (Suzhou) Co., Ltd Ethics Committee. The NHP study was designed and handled by Medicilon Preclinical Research (Shanghai) LLC, under the supervision of the IACUC of Medicilon Preclinical Research (Shanghai) LLC.

### Evaluation of *in vivo* delivery efficiency

To evaluate the delivery effect of LNP^Local^ and LNP^Lung^ *in vivo*, C57BL/6 mice aged 6-8 weeks were employed. We prepared luciferase mRNA-LNP for in vivo imaging in mice after 6 hours of injection. For the intramuscular (i.m.) injection of LNP^Local^, the mice’s legs were depilated one day in advance, followed by an i.m. injection the next day with a dosage of 0.05 mg/kg per leg. IVIS imaging was conducted after 6 hours of injection. For the intravenous (i.v.) injection of LNP^Lung^, the dosage was set at 0.25 mg/kg, and IVIS imaging was performed after 6 hours of injection. This experimental design allowed for the assessment of the *in vivo* delivery efficacy of LNP^Local^ and LNP^Lung^ formulations in the specified mouse model.

### Detection the concentration of IL-15 superagonists

The effective concentration of IL-15 in the samples was assessed using the Human IL-15 Quantikine ELISA Kit (R&D Systems, MN, USA). All subsequent steps after sample collections were carried out following the instructions provided by the kit. The actual concentration of the target molecules was determined by converting their molecular weight.

### Detection the binding affinity of IL-15 superagonists to IL-2Rβ/γ

To evaluate the binding affinity of the IL-15 superagonists to the IL-2Rβ/γ complex, affinity assays were conducted using standard ELISA methods. In brief, 96-well ELISA plates were coated by incubating them with 0.5 μg/mL of recombinant human IL-2Rβ/γ Heterodimer Protein at 4°C overnight. The plates were then washed three times with PBST and subsequently blocked with PBS containing 5% bovine serum albumin for 1 hour at 37°C. Following the washing of the plates, duplicate serial dilutions of mRNA supernatant or N-803 were added to the plates and incubated at room temperature for 2 hours. After this incubation, the plates were washed three times and treated with HRP-labeled Goat Anti-Human IgG-Fc Secondary Antibody at 37°C for 1 hour. Following further washing, the plates were exposed to a TMB single-component substrate solution and kept in the dark for 5-10 minutes. The reaction was stopped using 2 M sulfuric acid, and the absorbance was measured at both 450 nm and 570 nm. The final readout was determined using the formula: A=OD (450nm) – OD (570nm). All solutions used in this part, unless specified, were sourced from Multi Science (Hangzhou, China).

### Activation of IL-2R<u>β</u>-STAT5 signaling in TF-1 reporter cells

The IL-2Rβ-STAT5 reporter TF-1 cell line was engineered to possess stably expressed human IL-2Rβ on its cell membrane, along with a conditionally expressed luciferase module. This luciferase module is driven by a promoter region containing a STAT5 response element and a miniTATA box. The TF-1 cells were suspended in RPMI1640 medium supplemented with 10% FBS and 1% P/S, and were then seeded into a 96-well round bottom cell culture microplate (obtained from Corning, NY, USA) at a concentration of 1E5 cells per well. Subsequently, samples were added to the wells to reach a final volume of 200 μL per well. After co-culturing for 24 hours, the samples were collected for the assessment of luciferase activity using the One-Lite Luciferase Assay System (Vazyme, Nanjing, China). The luciferase activity was quantified using a plate-based luminometer (Tecan, Männedorf, Switzerland).

### Detection of IL-6 and IFN-<u>γ</u> level in PBMC activation assay

Peripheral blood mononuclear cells (PBMCs) were suspended in ImmunoCult™-XF T Cell Expansion Medium supplemented with 10 μg/ml CD3Ab (OKT3, obtained from Sino Biological, Beijing, China). Subsequently, the PBMCs were seeded into a 96-well round bottom cell culture microplate (obtained from Corning, NY, USA) at a concentration of 4E5 cells per well. Following this, samples were added to the wells to achieve a final volume of 200 μL per well. After co-culturing for a duration of 72 hours, the supernatant was collected for the assessment of IFN-**<u>γ</u>** levels using the Human IFN-gamma Quantikine ELISA Kit and IL6 levels using the Human IL-6 Quantikine ELISA Kit.

### Detection of pSTAT5 level and proliferation subclass in PBMC activation assay

Additionally, the cells were collected to evaluate the proliferation rate, the level of phosphorylated STAT5 (pSTAT5), and changes in the CD45+CD3+CD8+ and CD45+CD3+CD56+ cell subtypes. The proliferation rate was quantified using the CellTiter-Glo® Luminescent Cell Viability Assay (Promega, WI, USA), while the pSTAT5 level was assessed using the STAT5 alpha/beta (Phospho) [pY694/pY699] Human InstantOne™ ELISA Kit. For cell subtype analysis, the cells were washed and re-suspended in PBS containing 0.1% BSA, followed by staining with a panel of conjugated antibodies: anti-human CD45-APC-Cy7 antibody, anti-human CD3-FITC antibody, anti-human CD8-BV510 antibody, and anti-human CD56-PE-Cy7 antibody. Subsequently, the stained cells were washed twice to remove any unspecific binding antibodies and were then subjected to measurement using a Beckman Cytoflex S flow cytometer (Beckman, CA, USA). The resulting data were analyzed using FlowJo software (Tree Star; Ashland, OR, USA).

### Antitumor efficacy in mice

The comparative effectiveness of mRNA-encoded IL-15 superagonist and N-803 was evaluated in immunocompetent mice harboring subcutaneous (s.c.) tumors of B16F10 melanoma, CT26 colon carcinoma, or MC38 colon adenocarcinoma. Female C57BL/6 mice were subjected to subcutaneous implantation (s.c.) of tumor cell slurries consisting of B16F10 (1E6 cells/mouse) or MC38 (5E5 cells/mouse). Female Balb/c mice were subjected to subcutaneous implantation (s.c.) of tumor cell slurries consisting of CT26 (1E6 cells/mouse) cells. Tumor sizes were assessed using calipers, and tumor volumes were calculated using the formula: volume = length × width × width /2. For treatment administration, mRNAs were injected intratumorally in a fixed volume of 50 μL to mice with established tumors, while N-803 was administered intravenously. To assess the comparative efficacy of lung-targeted LNP, we employed immunocompetent mice with intravenous (i.v.) luciferase-expressing B16F10 melanoma. Female C57BL/6 mice were subjected to i.v. injection of B16F10 (5E5 cells/mouse) tumor cell slurries. Tumor growth was monitored through *in vivo* bioluminescence imaging, and six hours after tumor inoculation, mRNA was administered via i.v. injection in a consistent volume of 100 μL. Mice were euthanized when the tumor volume of any group reached 2,000 mm^3^, or when they exhibited signs of morbidity, such as hunched posture, ruffled fur, or reduced mobility, in accordance with the guidelines of the IACUC.

### Complete blood count and Comprehensive Metabolic Panel

Complete blood count (CBC) was conducted on whole blood samples using the Automatic Hematology Analyzer BC-5000 Vet (Mindray Bio-medical, Shenzhen, China). Comprehensive Metabolic Panel (CMP) was conducted on serum samples using the Hitachi Biochemical Analyzer 7100+ISE (Hitachi, Tokyo, Japan) with related reagents (FUJIFILM Wako Chemicals, Osaka, Japan).

### Analysis of CD8+ T cell and NK cell percentage in mice

The analysis of the percentage of CD8+ T cells and NK cells in mice were obtained by using flow cytometry. All agents mentioned in the figure were administered via intravenous injection to C57BL/6 mice. Seventy-two hours post-injection, the whole blood was collected, and PBMCs were isolated using a Mouse PBMC Isolation Kit. Subsequently, the cells were washed, suspended in PBS containing 0.1% BSA, and stained with a panel of fluorescence-conjugated antibodies: anti-mouse CD45-APC-Cy7 antibody, anti-mouse CD3-FITC antibody, anti-mouse CD8-PE-Cy7 antibody, and anti-mouse CD161-APC antibody. After staining, the cells were washed twice to eliminate any unspecific binding antibodies, and measurements were conducted using a Beckman Cytoflex S flow cytometer (Beckman, CA, USA). Data analysis was performed using FlowJo software (Tree Star; Ashland, OR, USA).

### Pharmacokinetics (PK)

This experimental design involved administering a single treatment to female C57BL/6 mice according to the agents outlined in the figures. Sample collections were conducted at various time points, including prior to dosing and at 4, 8, 24, 48, 72, 96, or 120 hours following a solitary dose injection. The administration of mRNAs was performed either intramuscularly using a consistent volume of 50 μL or intravenously using a consistent volume of 100 μL. Concurrently, N-803 was administered intravenously. The concentrations of the agents were determined using an ELISA kit, with the calculation methodology previously elucidated.

### Hematoxylin and Eosin staining

The initial steps for Hematoxylin and Eosin (H&E) staining involve the dehydration and paraffin embedding of lung tissue, followed by sectioning the tissue into slices with a thickness of 3 μm. Place the slices in an oven set to a temperature of 60 ℃ and allow them to bake for a duration of 30 minutes. The staining process involved the following steps: firstly, the sample was treated with xylene three times for durations of 10 minutes, 5 minutes, and 5 minutes respectively. This was followed by treatment with anhydrous ethanol twice, for durations of 5 minutes and 2 minutes. Subsequently, the sample was treated with 95% alcohol for 2 minutes, followed by 80% alcohol for 2 minutes. The sample was then washed with water for 2 minutes. Next, the sample was stained with hematoxylin staining solution for a period of 1-5 minutes, and subsequently washed with water for 5 minutes. Rapid differentiation of the differentiation solution was performed, followed by rinsing with warm water until the sample returned to a blue color. The sample was then stained with eosin staining solution for 5 minutes, followed by treatment with 95% alcohol for 1-2 minutes. Finally, the sample was treated with anhydrous ethanol three times for durations of 1-3 minutes each, and then with xylene twice for 2 minutes each time. Next, proceed to securely close and allow for natural evaporation of moisture. Ultimately, images were captured utilizing a microscope.

### Immunohistochemical staining

Immunohistochemical (IHC) staining was conducted on lung tissue sections by initially performing normal dewaxing, followed by a 30 minutes antigen retrieval process. Subsequently, the sections were washed three times with phosphate-buffered saline (PBS) for a duration of 5 minutes each. Subsequently, immunohistochemistry was conducted on the water blocking ring, which was rendered inactive by treating it with a 3% hydrogen peroxide solution for a duration of 10 minutes. Following this, the ring was subjected to three washes with PBS, each lasting for 5 minutes. Subsequently, a sealing solution was incrementally introduced over a period of 30 minutes, after which the sealing solution was subsequently drained. The initial antibody working solution was introduced directly and subjected to overnight incubation at a temperature of 4°C within a refrigerated environment. Subsequently, the sample was subjected to three consecutive washes with PBS, with each wash lasting for a duration of 5 minutes. Following this, the secondary antibody was subjected to incubation at a temperature of 37°C for a duration ranging from 10 to 30 minutes. Subsequently, it underwent three rounds of washing with PBS, with each wash lasting 5 minutes. Finally, horseradish peroxidase (HRP) was introduced. The samples were subjected to incubation at a temperature of 37°C for a duration of 10 to 30 minutes. Subsequently, they underwent three washes with PBS, with each wash lasting 5 minutes. Finally, the samples were subjected to DAB staining. It is recommended to perform a complete rinsing with tap water, followed by the subsequent steps of restaining, sealing, and observation under a fluorescence microscope.

### NHP *in vivo* studies

The NHP study was designed and handled by Medicilon Preclinical Research (Shanghai) LLC. The work was carried out in strict adherence to the Regulations for the Care and Use of Laboratory Animals and Guideline for Ethical Review of Animal Welfare (China, GB/T 35892-2018), under the supervision of the IACUC of Medicilon Preclinical Research (Shanghai) LLC. Three macaca fascicularis individuals, aged three to five years and weighing three to five kilogrammes, were used in NHP investigations. The macaca fascicularis (Non-naïve) were obtained from Medicillon’s animal reserve 999M-014. The experiment utilized Macaca fascicularis as the subjects, with a control group comprising of one female, and an experimental group consisting of one female and one male. The first intravenous administration of Macaca fasciculiris (LNP^Lung^ without Fluc, 0.25 mpk) occurred at 0 hours, followed by a second intravenous administration 24 hours later (LNP^Lung^ with Fluc, 0.125 mpk). Imaging was conducted 6 hours after the second injection to examine the presence of Fluc in different organs. In addition, frozen lung tissue slices were acquired to specifically investigate the expression of Fluc in the lungs. In addition, Comprehensive Metabolic Panel and Complete Blood Count were conducted.

### Data analysis

EC_50_ were calculated by Four Parameter Logistic (4PL) Regression. Binding affinity were assessed by receptor saturation one site specific binding Kd model. Comparisons of continuous variables were done using Student t tests or ANOVA (2-tailed). P values ≤ 0.05 were considered significant (*, P < 0.05; **, P < 0.01; ***, P < 0.001; ****, P < 0.0001). All calculation were executed by GraphPad Prism Version 8.0.2 (GraphPad Softeare, CA, USA). The graphical abstract image in the article was created using Biorender website (https://app.biorender.com).

## Acknowledgements

We thank employees of Starna Therapeutics for their helpful technical and scientific support with NHPs and materials. This study was supported in part from Starna Therapeutics (STR-P002), Jiangsu Provincial Science and Technology Project (SBK2023070025), Suzhou Science and Technology Project (ZXL2023267). We are very grateful to Professor Wei Tuo from the Institute of Zoology, Chinese Academy of Sciences for the support of Biorender.

## Author contributions

R.H. and Q.C. supervised the project. J.Y., Q.L., Q.C. and R.H. designed the experiments. J.Y., Q.L. and Q.C. wrote the manuscript. J.Y. and X.W. performed the entire molecular experiments.

Q.L. assisted in the preparation and characterization of LNP, and conducted part of mice experiments. S.L. assisted in the synthesis of ionizable cationic lipids. J.Y., Q.L., Q.C. and R.H. analyzed data. All the authors discussed the results and commentated manuscript.

## Supplementary Text

**Supplemental Figure S1.**
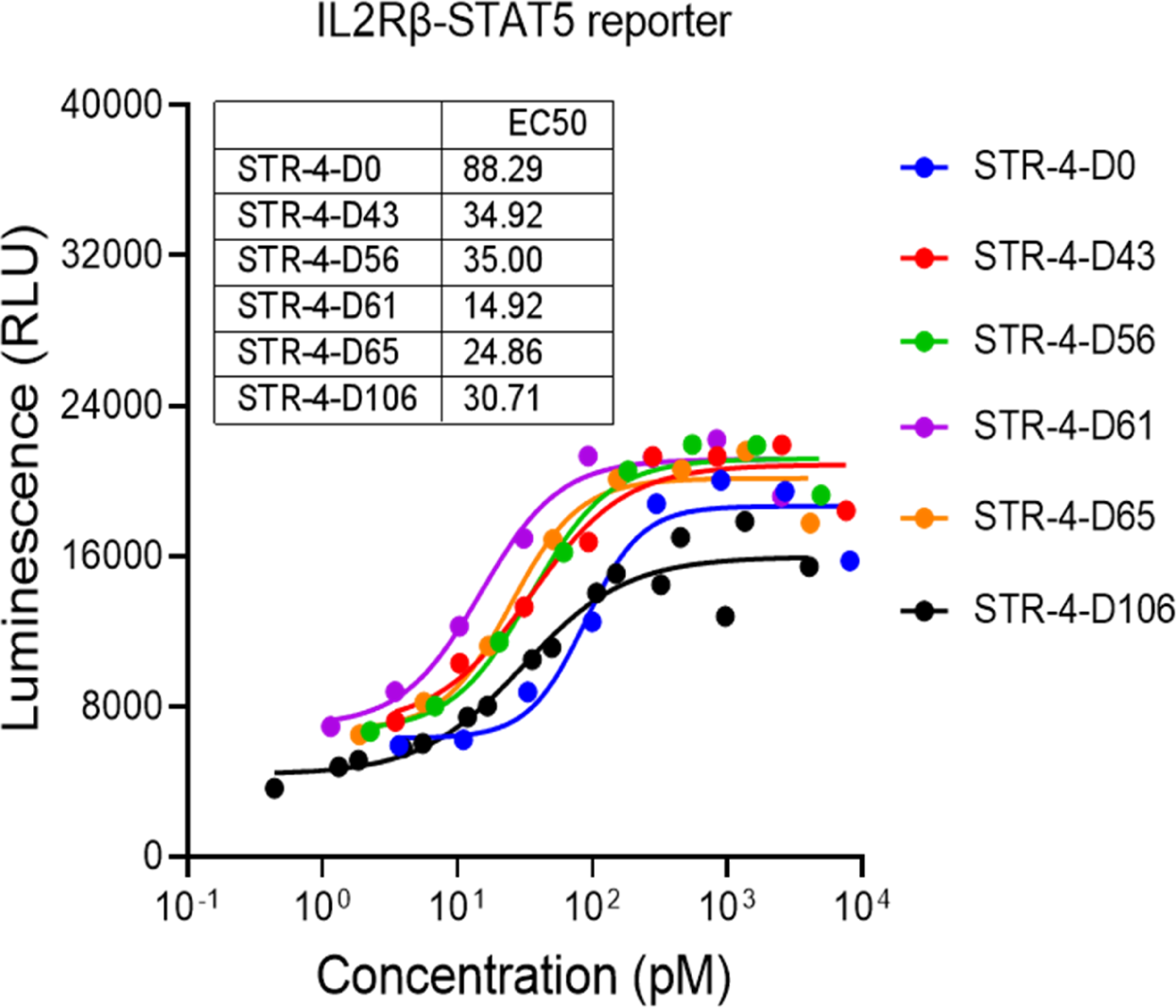
Detection of STR-4 variants inducing activating signal of IL2Rβ-STAT5 in TF-1 IL2Rβ-STAT5 reporter cells. Reporter cells were co-cultured with cell medium containing selected STR-4 ‘to D’ variants for 24 h before detection, presented by EC50 of inducing IL2Rβ-STAT5 reporter TF-1 signaling.

**Supplemental Figure S2.**
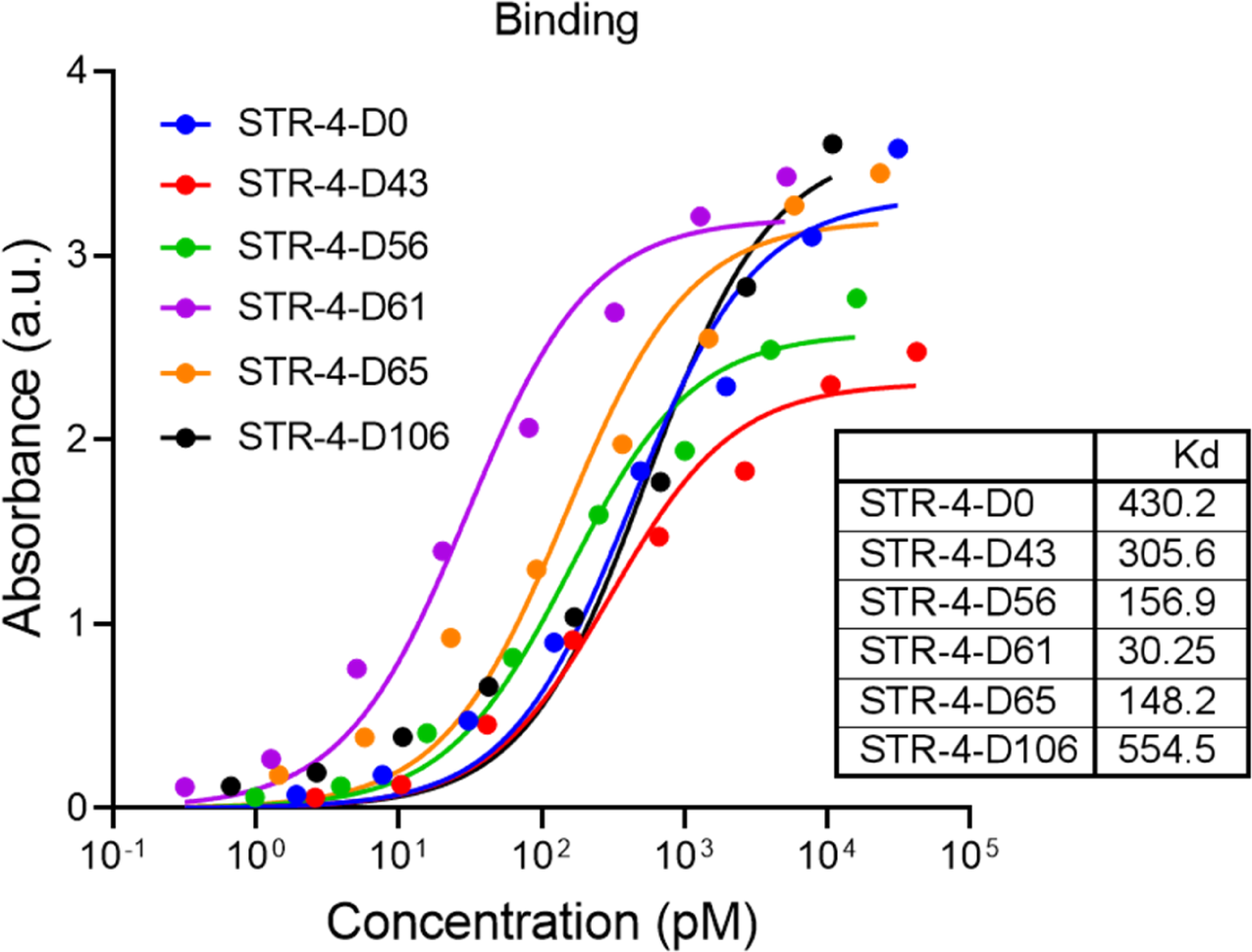
Detection of the binding affinity of STR-4 variants to IL-2Rβ/γ complex. Selected STR-4 ‘to D’ variants mRNAs were transfected to HEK293T cells through transfection reagent. Twenty-four hours after transfection, supernatants were collected to detect the potency. IL-2Rβ/γ complex protein were coated in plate, then selected STR-4 ‘to D’ variants supernatants were incubated overnight within the plate. Subsequently, Fc fragment of the variants were detected by 2^nd^ antibody to demonstrate the relative quantity of binding variants.

**Supplemental Figure S3.**
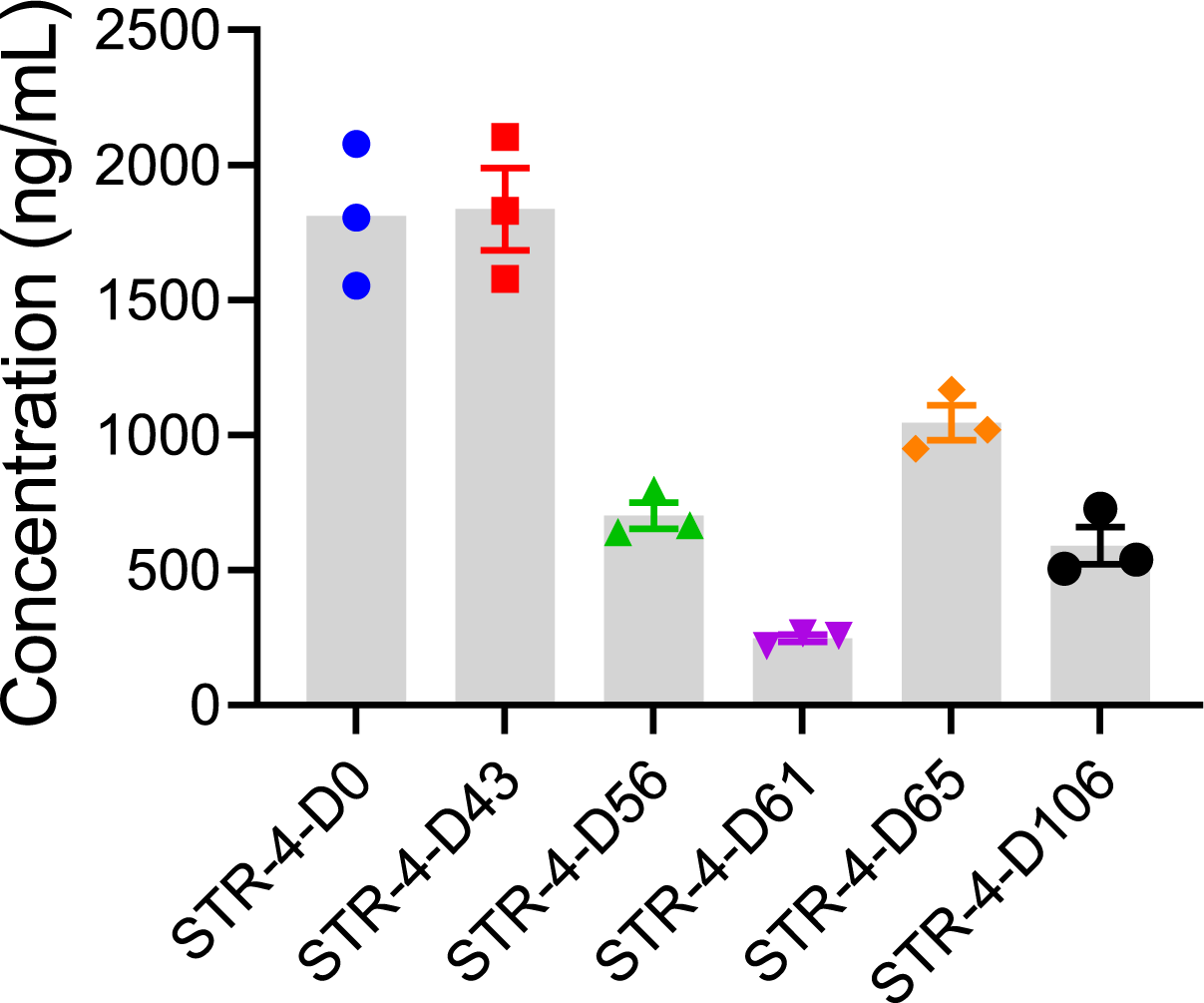
Expression level of selected STR-4 ‘to D’ variants in HEK293 cells. Selected STR-4 ‘to D’ variants mRNAs were transfected to HEK293T cells through transfection reagent. Twenty-four hours after transfection, supernatants were collected to detect the concentration of the variants.

**Supplemental Figure S4.**
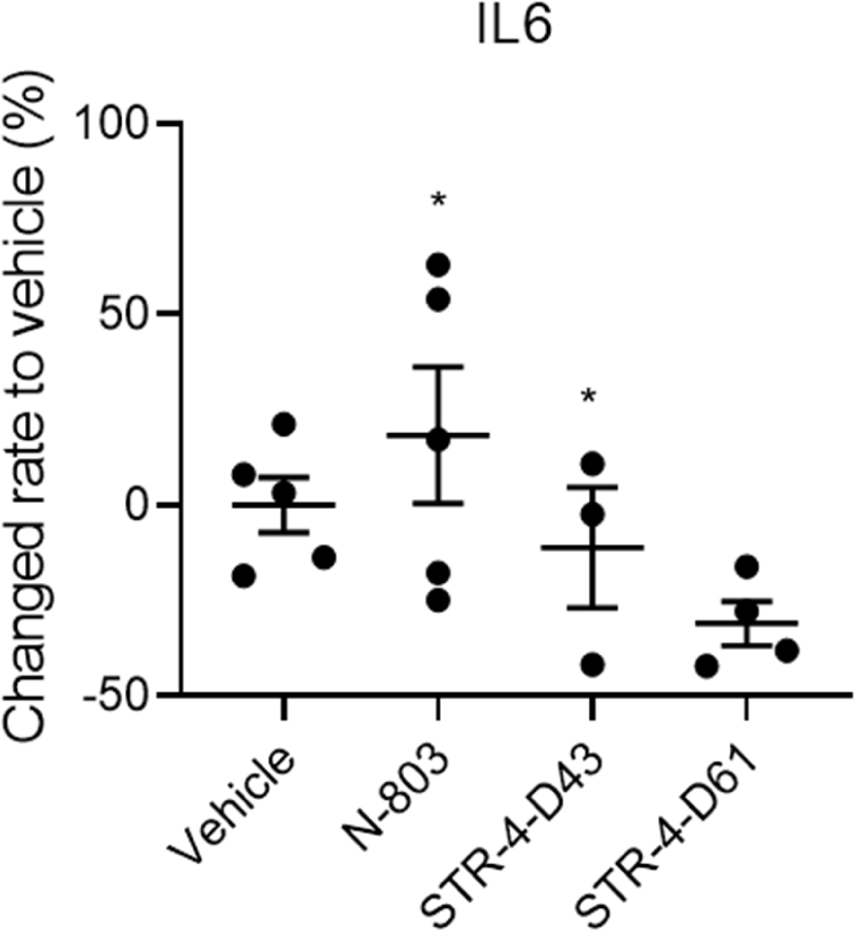
IL-6 level in MC38 tumor bearing mice after treatment of STR-4 variants. Blood samples were collected from MC38 tumor bearing mice on D17 after first dose, align with the schematic diagram from Fig.4F. Data is presented as mean ± standard error.

**Supplemental Figure S5.**
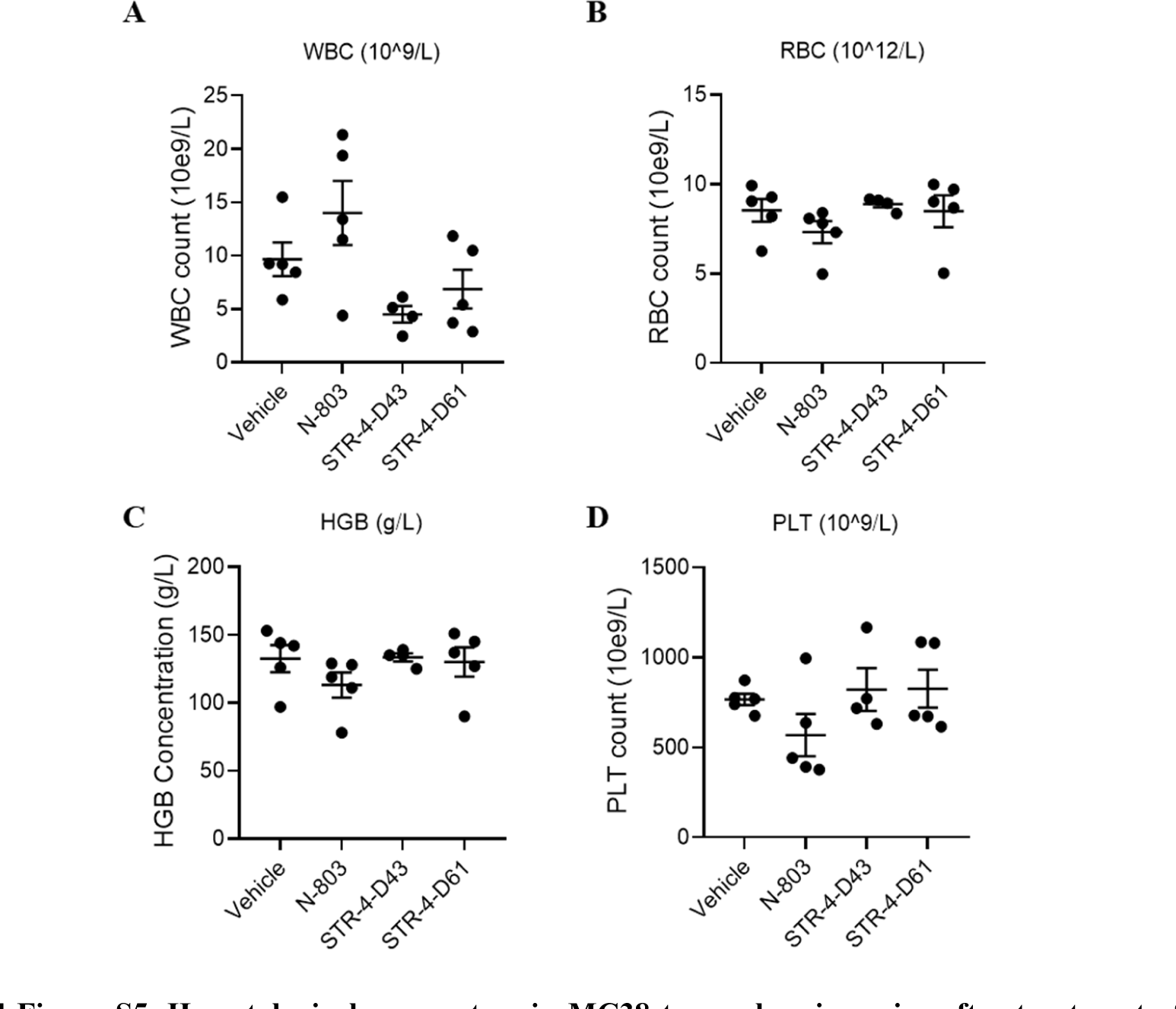
Hematological parameters in MC38 tumor bearing mice after treatment of STR-4 variants. Blood samples were collected from MC38 tumor bearing mice on D17 after first dose, align with the schematic diagram from Fig.4F. Samples were analyzed by Complete Blood Count within 6h after sampling. Data is presented as mean ± standard error.

**Supplemental Table 1.**
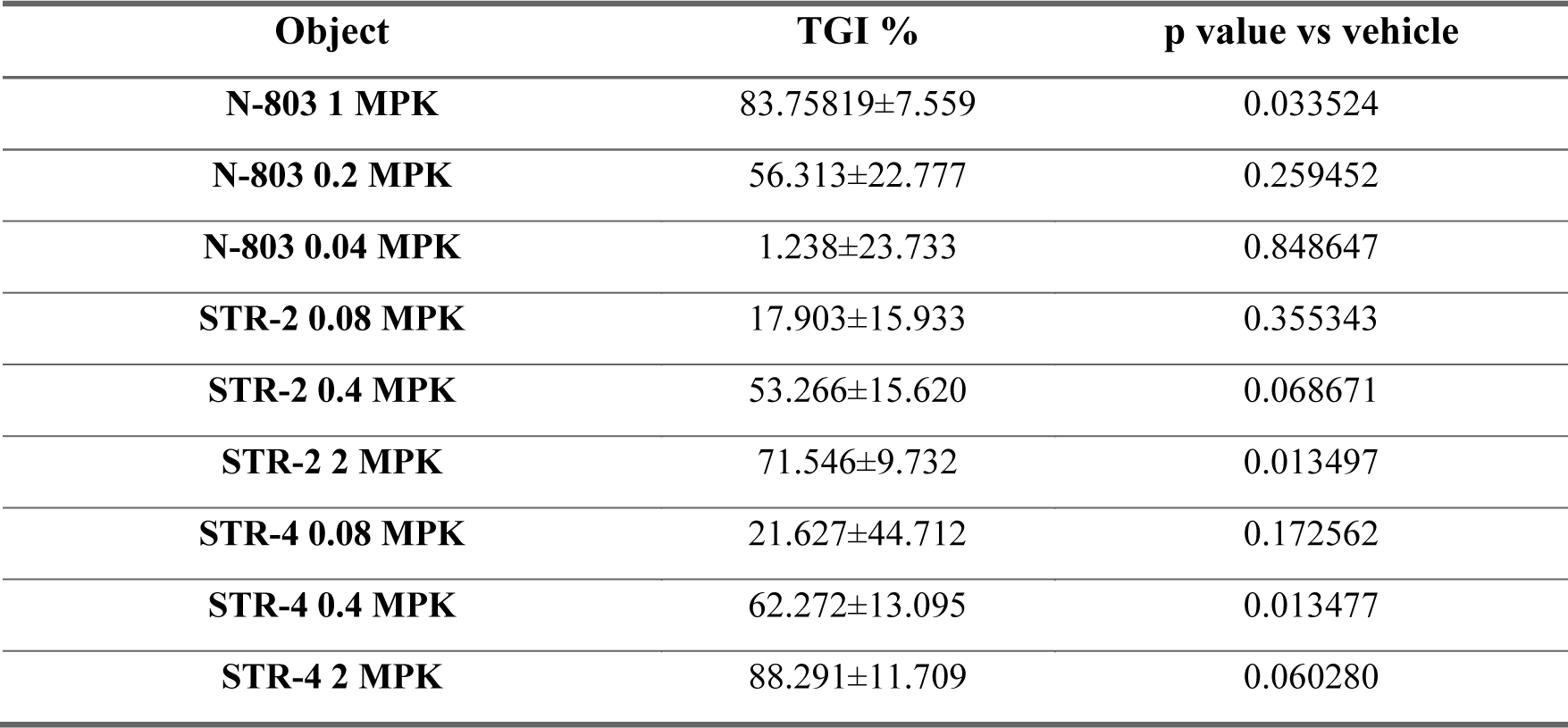
Tumor growth inhibition (TGI%) in CT26 tumor bearing mice. Tumor volume data were collected from CT26 tumor bearing mice on D12 after first dose, align with the schematic diagram from Fig.3A. TGI % were calculated by the equation of ‘[1 – (change of tumor volume in treatment group/change of tumor volume in control group)] × 100’. Data is presented as mean ± standard error.

**Supplemental Table 2.** Tumor growth inhibition (TGI%) in B16F10 tumor bearing mice. Tumor volume data were collected from B16F10 tumor bearing mice on D12 after first dose, align with the schematic diagram from Fig.3F.TGI % were calculated by the equation of ‘[1 – (change of tumor volume in treatment group/change of tumor volume in control group)] × 100’. Data is presented as mean ± standard error.

| object | TGI % | p value vs vehicle | p value vs N-803 0.2MPK |
| --- | --- | --- | --- |
| N-803 0.2MPK | 28.062±23.617 | 0.447881 | N.A. |
| STR-2 2MPK | 61.189±6.203 | 0.018652 | 0.122862 |
| STR-4 2MPK | 85.224±4.200 | 0.009325 | 0.038684 |

**Supplemental Table 3.** *In vitro* parameters of STR-4 ‘to D’ variants. STR-4 ‘to D’ variants mRNAs were transfected to HEK293T cells through transfection reagent. Twenty-four hours after transfection, supernatants were collected for *in vitro* characterization. Expression level was demonstrated by the concentration of variants in the supernatant. Affinity was evaluated by the binding affinity of the supernatants to IL-2Rβ/γ complex. EC50 of IL2Rβ-STAT5 signaling was measured by TF-1 reporter cells.

| Items | Expression<br>level (pg/ml) | Affinity to IL2Rbg complex |  | IL2R β -STAT5 signaling |
| --- | --- | --- | --- | --- |
|  |  | Bmax | Kd (nM) | EC50 (pM) |
| D1 | 21215.1 | 1.767 | 1869.808588 | 53.1491 |
| D2 | 27242.9 | 0.04885 | 4.424785689 | N.A. |
| D3 | 32887.7 | 3.21 | 4248.561475 | 53.58751 |
| D4 | 26330.0 | 1.937 | 2484.049008 | 72.35292 |
| D5 | 27743.0 | 0.03831 | 0.606803147 | N.A. |
| D6 | 6148.0 | 1.572 | 548.1661252 | N.A. |
| D7 | 33838.1 | 3.235 | 4219.908404 | 68.97875 |
| D8 | 21349.5 | 0.044 | 1.353583591 | 297.6475 |
| D9 | 14095.1 | 0.5264 | 806.591772 | 34.79861 |
| D10 | 19492.0 | 0.7883 | 1192.233922 | 43.23012 |
| D11 | 6815.3 | 0.03339 | 0.087055232 | N.A. |
| D12 | 19552.1 | 0.722 | 1110.345637 | 53.05515 |
| D13 | 10342.7 | 0.2161 | 260.2262497 | N.A. |
| D14 | 13098.7 | 0.3094 | 465.1818217 | 54.14334 |
| D15 | 17361.8 | 0.5951 | 966.6105609 | 92.6919 |
| D16 | 21409.2 | 0.9726 | 1267.780953 | 45.49262 |
| D17 | 14914.5 | 1.348 | 1268.328962 | 64.77473 |
| D18 | 31453.7 | 0.2735 | 1199.123185 | 23.14949 |
| D19 | 21215.1 | 0.8764 | 1619.44651 | 55.12976 |
| D20 | 32580.5 | 3.218 | 4301.561827 | 45.83708 |
| D21 | 16936.0 | 1.081 | 1424.35511 | 112.342 |
| D22 | 22348.1 | 0.2991 | 1036.520922 | 46.46338 |
| D23 | 36264.2 | 0.02645 | 0.467138999 | 24.86398 |
| D24 | 10230.1 | 0.07722 | 0.756957764 | 91.90903 |
| D25 | 17574.3 | 0.9299 | 781.8530552 | N.A. |
| D26 | 14667.6 | 0.6109 | 401.5344267 | N.A. |
| D27 | 18739.0 | 0.6731 | 716.091909 | N.A. |
| D28 | 30311.3 | 1.29 | 1355.932203 | 52.32708 |
| D29 | 18452.2 | 0.8087 | 758.7583669 | 80.71398 |
| D30 | 27404.7 | 1.178 | 1531.373547 | 66.55967 |
| D31 | 8471.3 | 0.05701 | 0.168708655 | 34.36803 |
| D32 | 24247.6 | 0.9508 | 1891.415822 | 63.06024 |
| D33 | 20422.4 | 1.157 | 1211.257682 | 43.05789 |
| D34 | 10133.5 | 24.64 | 12834.07054 | 56.40584 |
| D35 | 34262.1 | 0.711 | 1443.144009 | N.A. |
| D36 | 6165.3 | 0.009371 | 43.9581947 | 41.19466 |
| D37 | 5153.4 | 49.53 | 6608.134027 | 22.64062 |
| D38 | 11381.4 | 10.03 | 17521.27451 | N.A. |
| D39 | 9680.9 | N.A. | N.A. | 43.76248 |
| D40 | 15237.8 | 0.03563 | 85.56777704 | 32.12119 |
| D41 | 6815.3 | 0.01072 | 36.82624183 | 26.66458 |
| D42 | 34700.5 | 264.3 | 53149.88061 | N.A. |
| D43 | 27698.9 | N.A. | N.A. | 19.54045 |
| D44 | 5540.3 | N.A. | N.A. | 27.80757 |
| D45 | 35942.7 | N.A. | N.A. | 35.9416 |
| D46 | 11365.5 | N.A. | N.A. | 35.08044 |
| D47 | 5400.1 | 136.6 | 7165.22488 | 43.11269 |
| D48 | 5522.8 | 0.07082 | 38.77559009 | N.A. |
| D49 | 22838.8 | 0.02667 | 0.76697851 | 27.34568 |
| D50 | 8421.8 | 0.08987 | 0.980937096 | 49.01554 |
| D51 | 17998.6 | 1.03 | 1375.817121 | 33.50687 |
| D52 | 13660.2 | 1.304 | 1674.169178 | 87.75982 |
| D53 | 46806.5 | 2.753 | 2939.679806 | 46.12675 |
| D54 | 13722.4 | 0.2495 | 454.3782049 | 25.88171 |
| D55 | 10423.0 | 1.107 | 633.499041 | 36.20777 |
| D56 | 14822.0 | 2.053 | 1470.153051 | 16.68298 |
| D57 | 20437.4 | 0.1957 | 229.6160019 | 28.72353 |
| D58 | 22883.3 | 2.598 | 1559.948331 | 31.78455 |
| D59 | 46865.3 | 2.758 | 2351.274122 | N.A. |
| D60 | 14497.7 | 0.5473 | 423.2982346 | 75.57835 |
| D61 | 8273.1 | 2.327 | 469.5658981 | 14.57705 |
| D62 | 7356.0 | 1.844 | 497.1229499 | N.A. |
| D63 | 26049.9 | 1.263 | 1226.680236 | N.A. |
| D64 | 14048.6 | 0.5356 | 542.999178 | 54.41735 |
| D65 | 12848.3 | 1.182 | 417.5832779 | 14.02904 |
| D66 | 18935.0 | 0.7546 | 1032.998004 | 60.93083 |
| D67 | 26315.2 | 0.06229 | 1.721532861 | N.A. |
| D68 | 25016.7 | 0.9986 | 1172.662152 | 33.53818 |
| D69 | 12754.3 | 1.156 | 1206.795318 | 50.70654 |
| D70 | 24602.8 | 2.51 | 2026.461033 | 27.25956 |
| D71 | 7507.1 | 0.353 | 368.3407054 | 31.43226 |
| D72 | 23995.9 | 0.6037 | 190.7073238 | 29.06016 |
| D73 | 17149.0 | 1.477 | 1032.528281 | 33.82785 |
| D74 | 34905.2 | 2.495 | 1462.011195 | N.A. |
| D75 | 26226.8 | 1.824 | 1172.035855 | 28.51998 |
| D76 | 20152.6 | 1.206 | 905.3117783 | 43.73899 |
| D77 | 9323.4 | 0.3594 | 253.2586997 | 50.01761 |
| D78 | 16448.3 | 0.9087 | 795.631581 | 46.51035 |
| **D79** | 20512.2 | 0.03858 | -<br>0.373820801 | 41.18683 |
| **D80** | 19972.6 | 0.5469 | 622.53885 | 38.91651 |
| **D81** | 11651.1 | 0.124 | 124.0067327 | N.A. |
| **D82** | 37930.1 | 0.04284 | 2.222570165 | 25.20061 |
| **D83** | 19552.1 | 0.02986 | 0.361060007 | 51.02752 |
| **D84** | 31556.2 | 0.0496 | 18.0999726 | 32.76314 |
| **D85** | 37170.2 | 0.03827 | 13.12091439 | 33.92179 |
| **D86** | 18512.6 | 0.7136 | 808.7838102 | 51.14495 |
| **D87** | 26521.5 | 0.4821 | 1329.784319 | 32.26211 |
| **D88** | 10052.8 | 0.03902 | 5.262457431 | 65.18182 |
| **D89** | 11888.5 | 0.1838 | 338.9047638 | 74.95988 |
| **D90** | 10101.2 | 0.07921 | 67.89055466 | 56.23361 |
| **D91** | 14791.1 | 0.4102 | 340.3922183 | 51.16843 |
| **D92** | 29006.1 | 1.034 | 1263.710025 | 32.87274 |
| **D93** | 14404.9 | 0.0212 | 0.128860532 | 54.68352 |
| **D94** | 24336.4 | 0.02101 | 0.208321917 | 33.80436 |
| **D95** | 14729.4 | 0.4672 | 713.117 | 56.27275 |
| **D96** | 29519.6 | 0.4432 | 1259.325948 | 37.87529 |
| **D97** | 18467.3 | 0.01794 | -<br>0.020808706 | 71.77359 |
| **D98** | 7187.6 | 0.307 | 148.5105883 | N.A. |
| **D99** | 6251.3 | 0.2226 | 129.9565507 | N.A. |
| **D100** | 27463.6 | 0.03528 | 0.312835167 | 50.80049 |
| **D101** | 25341.7 | 0.03482 | 0.26155713 | 76.64305 |
| **D102** | 4993.9 | 1.591 | 792.7349591 | N.A. |
| **D103** | 38967.8 | 5.15 | 8175.441343 | 34.72032 |
| **D104** | 22258.8 | 0.1269 | 299.7612244 | 49.11731 |
| **D105** | 38543.9 | 1.891 | 3669.080518 | 49.54789 |
| **D106** | 17255.4 | 1.748 | 915.2542373 | 17.57545 |
| **D107** | 43020.8 | 1.056 | 1791.286648 | 45.06987 |
| **D108** | 23936.6 | 0.6149 | 870.8654637 | 47.49677 |

## References

1. T. A. Fehniger, M. A. Caligiuri, Interleukin 15: biology and relevance to human disease. Blood. 97, 14–32 (2001).

2. K. Yoshihara, T. Yajima, C. Kubo, Y. Yoshikai, Role of interleukin 15 in colitis induced by dextran sulphate sodium in mice. Gut 55, 334–341 (2006).

3. M. Desbois, C. Béal, M. Charrier, B. Besse, G. Meurice, N. Cagnard, Y. Jacques, D. Béchard, L. Cassard, N. Chaput, IL-15 superagonist RLI has potent immunostimulatory properties on NK cells: Implications for antimetastatic treatment. J. ImmunoTher. Cancer 8, (2020).

4. K. M. Knudson, J. W. Hodge, J. Schlom, S. R. Gameiro, Rationale for IL-15 superagonists in cancer immunotherapy. Expert Opin. Biol. Ther. 20, 705–709 (2020).

5. J. C. Steel, T. A. Waldmann, J. C. Morris, Interleukin-15 biology and its therapeutic implications in cancer. Trends Pharmacol. Sci. 33, 35–41 (2012).

6. W. Chen, N. Liu, Y. Yuan, M. Zhu, X. Hu, W. Hu, S. Wang, C. Wang, B. Huang, D. Xing, ALT-803 in the treatment of non-muscle-invasive bladder cancer: Preclinical and clinical evidence and translational potential. Front. Immunol. 13, 1040669 (2022).

7. H. Furuya, O. Chan, I. Pagano, C. Zhu, N. Kim, R. Peres, K. Hokutan, S. Alter, P. Rhode, C. J. Rosser, Effectiveness of two different dose administration regimens of an IL-15 superagonist complex (ALT-803) in an orthotopic bladder cancer mouse model. J. Transl. Med. 17, 1–12 (2019).

8. M. Ahdoot, D. Theodorescu, Immunotherapy of high risk non-muscle invasive bladder cancer. Expert Rev. Clin. Pharmacol. 14, 1345–1352 (2021).

9. G. Gakis, Adjuvant instillation therapy for non-muscle invasive bladder cancer-beyond BCG und mitomycin C. Aktuelle Urologie, (2022).

10. R. Romee, S. Cooley, M. M. Berrien-Elliott, P. Westervelt, M. R. Verneris, J. E. Wagner, D. J. Weisdorf, B. R. Blazar, C. Ustun, T. E. DeFor, First-in-human phase 1 clinical study of the IL-15 superagonist complex ALT-803 to treat relapse after transplantation. Blood. 131, 2515–2527 (2018).

11. Y. Peng, S. Fu, Q. Zhao, 2022 update on the scientific premise and clinical trials for IL-15 agonists as cancer immunotherapy. J. Leukocyte Biol. 112, 823–834 (2022).

12. K. Margolin, C. Morishima, V. Velcheti, J. S. Miller, S. M. Lee, A. W. Silk, S. G. Holtan, A. M. Lacroix, S. P. Fling, J. C. Kaiser, Phase I trial of ALT-803, a novel recombinant IL15 complex, in patients with advanced solid tumors. Clin. Cancer Res. 24, 5552–5561 (2018).

13. Y. Guo, L. Luan, N. K. Patil, E. R. Sherwood, Immunobiology of the IL-15/IL-15Rα complex as an antitumor and antiviral agent. Cytokine Growth Factor Rev. 38, 10–21 (2017).

14. K. M. Knudson, K. C. Hicks, Y. Ozawa, J. Schlom, S. R. Gameiro, Functional and mechanistic advantage of the use of a bifunctional anti-PD-L1/IL-15 superagonist. J. ImmunoTher. Cancer 8, (2020).

15. K. M. Knudson, K. C. Hicks, S. Alter, J. Schlom, S. R. Gameiro, Mechanisms involved in IL-15 superagonist enhancement of anti-PD-L1 therapy. J. ImmunoTher. Cancer 7, 1–16 (2019).

16. T. A. Waldmann, in J. Investig. Dermatol. Symp. Proc. (Elsevier, 2013), vol. 16, pp. S28–S30.

17. M. T. Williams, Y. Yousafzai, C. Cox, A. Blair, R. Carmody, S. Sai, K. E. Chapman, R. McAndrew, A. Thomas, A. Spence, Interleukin-15 enhances cellular proliferation and upregulates CNS homing molecules in pre-B acute lymphoblastic leukemia. Blood. 123, 3116–3127 (2014).

18. J. Chen, M. Petrus, R. Bamford, J. H. Shih, J. C. Morris, J. E. Janik, T. A. Waldmann, Increased serum soluble IL-15Rα levels in T-cell large granular lymphocyte leukemia. Blood. 119, 137–143 (2012).

19. R. Zhang, M. V. Shah, J. Yang, S. B. Nyland, X. Liu, J. K. Yun, R. Albert, T. P. Loughran Jr, Network model of survival signaling in large granular lymphocyte leukemia. Proc. Natl. Acad. Sci. U.S.A. 105, 16308–16313 (2008).

20. R. Zambello, M. Facco, L. Trentin, R. Sancetta, C. Tassinari, A. Perin, A. Milani, G. Pizzolo, F. Rodeghiero, C. Agostini, Interleukin-15 triggers the proliferation and cytotoxicity of granular lymphocytes in patients with lymphoproliferative disease of granular lymphocytes. Blood. 89, 201–211 (1997).

21. M. K. Kennedy, M. Glaccum, S. N. Brown, E. A. Butz, J. L. Viney, M. Embers, N. Matsuki, K. Charrier, L. Sedger, C. R. Willis, Reversible defects in natural killer and memory CD8 T cell lineages in interleukin 15–deficient mice. J. Exp. Med. 191, 771–780 (2000).

22. J. P. Lodolce, P. R. Burkett, D. L. Boone, M. Chien, A. Ma, T cell–independent interleukin 15rα signals are required for bystander proliferation. J. Exp. Med. 194, 1187–1194 (2001).

23. T. A. Waldmann, E. Lugli, M. Roederer, L. P. Perera, J. V. Smedley, R. P. Macallister, C. K. Goldman, B. R. Bryant, J. M. Decker, T. A. Fleisher, Safety (toxicity), pharmacokinetics, immunogenicity, and impact on elements of the normal immune system of recombinant human IL-15 in rhesus macaques. Blood. 117, 4787–4795 (2011).

24. C. Berger, M. Berger, R. C. Hackman, M. Gough, C. Elliott, M. C. Jensen, S. R. Riddell, Safety and immunologic effects of IL-15 administration in nonhuman primates. Blood. 114, 2417–2426 (2009).

25. M. C. Sneller, W. C. Kopp, K. J. Engelke, J. L. Yovandich, S. P. Creekmore, T. A. Waldmann, H. C. Lane, IL-15 administered by continuous infusion to rhesus macaques induces massive expansion of CD8+ T effector memory population in peripheral blood. Blood. 118, 6845–6848 (2011).

26. E. Lugli, C. K. Goldman, L. P. Perera, J. Smedley, R. Pung, J. L. Yovandich, S. P. Creekmore, T. A. Waldmann, M. Roederer, Transient and persistent effects of IL-15 on lymphocyte homeostasis in nonhuman primates. Blood. 116, 3238–3248 (2010).

27. ImmunityBio. 2022. “ImmunityBio Announces FDA Acceptance of Biologics License Application for N-803 in BCG-Unresponsive Non-Muscle-Invasive Bladder Cancer Carcinoma In Situ” ImmunityBio Official Website, June 28, 2022. https://immunitybio.com/immunitybio-announces-fda-acceptance-of-biologics-license-application-for-n-803-in-bcg-unresponsive-non-muscle-invasive-bladder-cancer-carcinoma-in-situ/.

28. M. Gaviria, B. Kilic, A network analysis of COVID-19 mRNA vaccine patents. Nat. Biotechnol. 39, 546–548 (2021).

29. Y.-K. Kim, RNA therapy: rich history, various applications and unlimited future prospects. Exp. Mol. Med. 54, 455–465 (2022).

30. Y. Weng, C. Li, T. Yang, B. Hu, M. Zhang, S. Guo, H. Xiao, X.-J. Liang, Y. Huang, The challenge and prospect of mRNA therapeutics landscape. Biotechnol. Adv. 40, 107534 (2020).

31. S. Chen, X. Huang, Y. Xue, E. Álvarez-Benedicto, Y. Shi, W. Chen, S. Koo, D. J. Siegwart, Y. Dong, W. Tao, Nanotechnology-based mRNA vaccines. Nat. Rev. Methods Primers 3, 63 (2023).

32. Q. Chen, Y. Zhang, H. Yin, Recent advances in chemical modifications of guide RNA, mRNA and donor template for CRISPR-mediated genome editing. Adv. Drug Delivery Rev. 168, 246–258 (2021).

33. X. Huang, N. Kong, X. Zhang, Y. Cao, R. Langer, W. Tao, The landscape of mRNA nanomedicine. Nat. Med. 28, 2273–2287 (2022).

34. Y. Zong, Y. Lin, T. Wei, Q. Cheng, Lipid Nanoparticle (LNP) Enables mRNA Delivery for Cancer Therapy. Adv. Mater., 2303261 (2023).

35. E. Kon, N. Ad-El, I. Hazan-Halevy, L. Stotsky-Oterin, D. Peer, Targeting cancer with mRNA–lipid nanoparticles: key considerations and future prospects. Nat. Rev. Clin. Oncol., 1–16 (2023).

36. C. Liu, Q. Shi, X. Huang, S. Koo, N. Kong, W. Tao, mRNA-based cancer therapeutics. Nat. Rev. Cancer, 1–18 (2023).

37. S. H. Kiaie, N. Majidi Zolbanin, A. Ahmadi, R. Bagherifar, H. Valizadeh, F. Kashanchi, R. Jafari, Recent advances in mRNA-LNP therapeutics: immunological and pharmacological aspects. J. Nanobiotechnol. 20, 276 (2022).

38. L. Miao, L. Li, Y. Huang, D. Delcassian, J. Chahal, J. Han, Y. Shi, K. Sadtler, W. Gao, J. Lin, J. C. Doloff, R. Langer, D. G. Anderson, Delivery of mRNA vaccines with heterocyclic lipids increases anti-tumor efficacy by STING-mediated immune cell activation. Nat. Biotechnol. 37, 1174–1185 (2019).

39. T. Wei, Q. Cheng, L. Farbiak, D. G. Anderson, R. Langer, D. J. Siegwart, Delivery of Tissue-Targeted Scalpels: Opportunities and Challenges for In Vivo CRISPR/Cas-Based Genome Editing. ACS Nano 14, 9243–9262 (2020).

40. K. L. Swingle, H. C. Safford, H. C. Geisler, A. G. Hamilton, A. S. Thatte, M. M. Billingsley, R. A. Joseph, K. Mrksich, M. S. Padilla, A. A. Ghalsasi, Ionizable lipid nanoparticles for in vivo mRNA delivery to the placenta during pregnancy. J. Am. Chem. Soc. 145, 4691–4706 (2023).

41. Q. Cheng, T. Wei, L. Farbiak, L. T. Johnson, S. A. Dilliard, D. J. Siegwart, Selective organ targeting (SORT) nanoparticles for tissue-specific mRNA delivery and CRISPR–Cas gene editing. Nat. Nanotechnol. 15, 313–320 (2020).

42. K.-p. Han, X. Zhu, B. Liu, E. Jeng, L. Kong, J. L. Yovandich, V. V. Vyas, W. D. Marcus, P.-A. Chavaillaz, C. A. Romero, IL-15: IL-15 receptor alpha superagonist complex: high-level co-expression in recombinant mammalian cells, purification and characterization. Cytokine 56, 804–810 (2011).

43. T. A. Waldmann, The biology of interleukin-2 and interleukin-15: implications for cancer therapy and vaccine design. Nat. Rev. Immunol. 6, 595–601 (2006).

44. X. Zhu, W. D. Marcus, W. Xu, H.-i. Lee, K. Han, J. O. Egan, J. L. Yovandich, P. R. Rhode, H. C. Wong, Novel human interleukin-15 agonists. J. Immunol. 183, 3598–3607 (2009).

45. M. Cai, X. Huang, X. Huang, D. Ju, Y. Z. Zhu, L. Ye, Research progress of interleukin-15 in cancer immunotherapy. Front. Pharmacol. 14, 1184703 (2023).

46. Y. Hailemichael, D. H. Johnson, N. Abdel-Wahab, W. C. Foo, S.-E. Bentebibel, M. Daher, C. Haymaker, K. Wani, C. Saberian, D. Ogata, Interleukin-6 blockade abrogates immunotherapy toxicity and promotes tumor immunity. Cancer Cell 40, 509–523. e506 (2022).

47. M. Wang, X. Zhai, J. Li, J. Guan, S. Xu, Y. Li, H. Zhu, The role of cytokines in predicting the response and adverse events related to immune checkpoint inhibitors. Front. Immunol. 12, 2894 (2021).

48. A. V. Hirayama, C. K. Chou, T. Miyazaki, R. N. Steinmetz, H. A. Di, S. P. Fraessle, J. Gauthier, S. Fiorenza, R. M. Hawkins, W. W. Overwijk, A novel polymer-conjugated human IL-15 improves efficacy of CD19-targeted CAR T-cell immunotherapy. Blood Adv. 7, 2479–2493 (2023).

49. T. A. Waldmann, S. Dubois, M. D. Miljkovic, K. C. Conlon, IL-15 in the combination immunotherapy of cancer. Front. Immunol. 11, 868 (2020).

50. Y. Yang, A. Lundqvist, Immunomodulatory Effects of IL-2 and IL-15; Implications for Cancer Immunotherapy. Cancers 12, 3586 (2020).

51. Y. Zhou, T. Husman, X. Cen, T. Tsao, J. Brown, A. Bajpai, M. Li, K. Zhou, L. Yang, Interleukin 15 in cell-based cancer immunotherapy. Int. J. Mol. Sci. 23, 7311 (2022).

